# Indirect pathway neurons in the tail of the striatum regulate inhibitory control over sensory driven behavior

**DOI:** 10.1101/2025.08.15.670612

**Authors:** Sarah M. Ferrigno, Nathan Zhang, Evan Iliakis, Saurabh Pandey, Jamie Galanaugh, Marc V. Fuccillo

## Abstract

Inhibitory control, or the ability to withhold action in certain situations, is behaviorally essential. Disrupted inhibitory control is linked to various neuropsychiatric symptoms, making it critical to understand the underlying neural basis. We examined how the tail of the striatum (TS), a major basal ganglia sensory hub, regulates actions to sensory stimuli. Mice performed an auditory Go/NoGo task where we recorded cell-specific activity of TS neurons. Both major striatal types were active during target sounds, but non-target sounds preferentially engaged indirect pathway neurons. Temporarily silencing this activity increased errors to non-target stimuli, indicating a role in suppressing inappropriate action. In mice deficient for the synaptic adhesion molecule Neurexin1α, a gene linked to autism spectrum disorder and ADHD, TS indirect pathway recruitment was reduced, and these mice demonstrated auditory-specific inhibitory control deficits. Altogether, these findings highlight a subcortical target to potentially improve attentional and behavioral regulation in neurodevelopmental disorders.

**Teaser:** Posterior striatal circuits control sensory-guided actions and are disrupted in a rodent model of neurodevelopmental disorders.

## Introduction

Inhibitory control, or the ability to suppress inappropriate urges or actions depending on situational context, is an essential part of daily function. Disruptions to this process likely drive impulsivity and attention deficits found in many neurodevelopmental disorders (NDDs). Traditional models have typically framed inhibitory control in terms of two distinct mechanisms — reactive inhibitory control, thought to be driven by bottom-up responses to unexpected sensory input, and proactive control, guided by top-down restraint in alignment with current goals^1–4^. However, these binary frameworks have failed to fully align with emerging evidence, and more recent proposals have conceptualized inhibitory control as a complex, continuous process shaped by the interaction of multiple mechanisms across broad neural networks^5–7^. In line with this, the classical view of inhibitory control as being primarily mediated by frontal executive circuits has expanded to include the basal ganglia (BG), whose established role in goal-directed behavior suggests they could serve as critical subcortical contributors to these cognitive processes^1,2,8^.

The BG are an evolutionarily conserved set of sub-cortical nuclei that can select and amplify cortical activity while simultaneously regulating midbrain targets^9,10^. While early experiments focused on the role of BG stimulation in generating movement^11,12^, recent work has revealed a broader involvement in regulating behavior. As the primary input structure of the BG, the striatum receives topographically organized projections from the entire neocortex, making it well suited to integrate goal-directed strategies and sensory context with action selection^13,14^. Cortical signals are processed in the striatum primarily through recruitment of its main neuronal population, the spiny projection neurons (SPNs)^15^. These neurons comprise roughly 95% of striatal neurons and consist of two equally sized subtypes with opposing effects on basal ganglia output^16–18^. Dopamine D1 receptor-expressing direct pathway SPNs (dSPNs) and D2 receptor-expressing indirect pathway SPNs (iSPNs) are thought promote respective excitation and inhibition of downstream thalamic neurons through distinct projections to intermediary BG nuclei^18^.

Given that iSPNs suppress thalamic output in part through polysynaptic disinhibition of the sub-thalamic nucleus (STN), a region previously implicated with inhibitory control, the indirect pathway is likely a key circuit element regulating control over inappropriate action^1,2^. Indeed, iSPNs have been implicated in promoting a range of inhibitory motor and non-motor avoidance behaviors^19–23^. In addition to its downstream connectivity, the role of iSPN-mediated lateral inhibition within striatum has gained increased attention^21,24–26^. Recent work has revealed that optogenetic activation of iSPNs in the dorsomedial striatum induces motor slowing through a pathway distinct from the downstream pallidal effects on transient punishment^21^. Together this suggests that iSPNs can regulate inhibitory control over actions through multiple means, making them a compelling candidate for further investigation.

As the striatum receives widespread input, understanding how iSPNs contribute to sensory associated inhibitory control requires identification of the relevant striatal subregions integrating sensory input with higher-order goals. While there is an established literature in perceptual decision-making within anterior striatal circuits^27–31^, recent work has identified the posterior tail of the striatum (TS) as an additional multi-modal sensory integration domain^32–36^. In line with this, several studies have shown that pathway-specific manipulation of SPNs within the TS is sufficient to bias lateral auditory choices in two-alternative forced-choice (2-AFC) behavioral tasks^26,37,38^. In addition to auditory processing, the TS has also been implicated in modulation of avoidance responses to threatening stimuli, and recent work has identified contributions of TS dSPNs and iSPNs for promoting or overcoming threat avoidance, respectively^39^. Taken together, it seems possible that the iSPNs in the TS serve as mediators of inhibitory control across different situational contexts. Indeed, findings in rhesus monkeys suggest the indirect pathway is involved in the rejection of low-value visual targets^40,41^. However, to date, TS iSPN involvement in sensory-driven inhibitory control has not been investigated at a cellular level.

While a growing body of work has implicated striatal synaptic dysregulation in genetic models of NDDs^17,42–44^, the specific contributions to disease-relevant behavioral changes remains understudied. Specifically, the potential role of iSPN dysfunction in mediating the impulsivity and attentional perturbations of NDDs is unknown. Recent work in slice has shown that mice lacking either one or both neurexin1α (Nrxn1α) alleles exhibit synaptic deficits for PFC connections onto iSPNs in the dorsomedial striatum^45^. Neurexins are presynaptic cell-adhesion molecules that regulate the development and maintenance of synapses throughout the brain^46–48^. Copy number variations impacting the Nrxn1α isoform have been associated with increased risk for schizophrenia, autism spectrum disorder (ASD), attention deficit hyperactivity disorder, and Tourette’s Syndrome^49–56^. Furthermore, Nrxn1α knockout mice likely have impaired sensory-related inhibitory control, manifest as reduced pre-pulse inhibition^57^. However, whether these striatal circuit dysfunctions contribute to sensorimotor impulsivity seen in NDDs remains to be directly assessed.

To investigate the underlying circuits and molecular mediators of inhibitory control, we designed a lick-based Go/NoGo task where lick responses to distinct bandlimited sounds were either rewarded or punished with a timeout. We recorded pathway-specific SPN recruitment via fiber photometry as mice performed this auditory-guided task. While on Go trials we find similar absolute peak fluorescence across both SPN subtypes, we find greater iSPN activity on NoGo trials, suggesting iSPNs in the tail of the striatum (TS) may play a role in suppressing responses to inappropriate auditory stimuli. Consistent with this, we found that optogenetic inhibition of iSPNs but not dSPNs is sufficient to increase erroneous false alarm responses. Interestingly, we also observed decreased excitatory synaptic recruitment of TS iSPNs in Nrxn1α heterozygotes, revealing a disruption in TS iSPN circuits relevant for associated NDDs. To assess whether this circuit dysfunction also accompanied impaired inhibitory control over sound driven behavior, we trained Nrxn1α knockout mice in our task and found that they exhibited specific increases in false alarm errors that could not be explained by general motor impulsivity. Altogether these data suggest that iSPN activity in the TS supports inhibitory control over auditory driven actions. Furthermore, because Nrxn1α mutations have been linked with various NDDs, our findings provide important insight into a subcortical circuit for inhibitory control that may provide unique therapeutic access for attentional and behavioral dysregulation.

## Results

To investigate mechanisms of inhibitory control, we employed an auditory Go/NoGo task in which mice were trained to appropriately respond or withhold responding to sounds associated with distinct outcomes (Fig. 1A, B). Mice were head-fixed in a custom-made behavioral apparatus where lick responses were recorded by an infrared optical lickometer at the reward spout. A speaker delivered bandlimited auditory stimuli, which were assigned as either “Go” or “NoGo” sounds. During Go trials, if mice licked within 1.15s of the target sound onset, the trial was considered a “hit” and animals received a 4uL water reward. Conversely, if animals failed to register a lick response within the response window following Go sound presentation, that trial was classified as a “miss” and the mice received a 5s lockout penalty. For NoGo trials, if an animal followed the non-target sound with a lick response, that trial was classified as a “false alarm,” resulting in a 5s lockout penalty. However, if the mouse correctly withheld responses after the NoGo sound, the trial was a “correct rejection,” which was unrewarded but bypassed the lockout. We used established signal detection theory metrics, including the discrimination index (d′) to quantify the ability to distinguish target and non-target stimuli, calculated as the difference of the z-transformed hit and false alarm rates, and the response bias (c) to measure the subject’s overall tendency to favor one response over the other, independent of the sound properties (calculated as the negative average of the z-transformed hit rate and false alarm rate).

**Figure 1.**
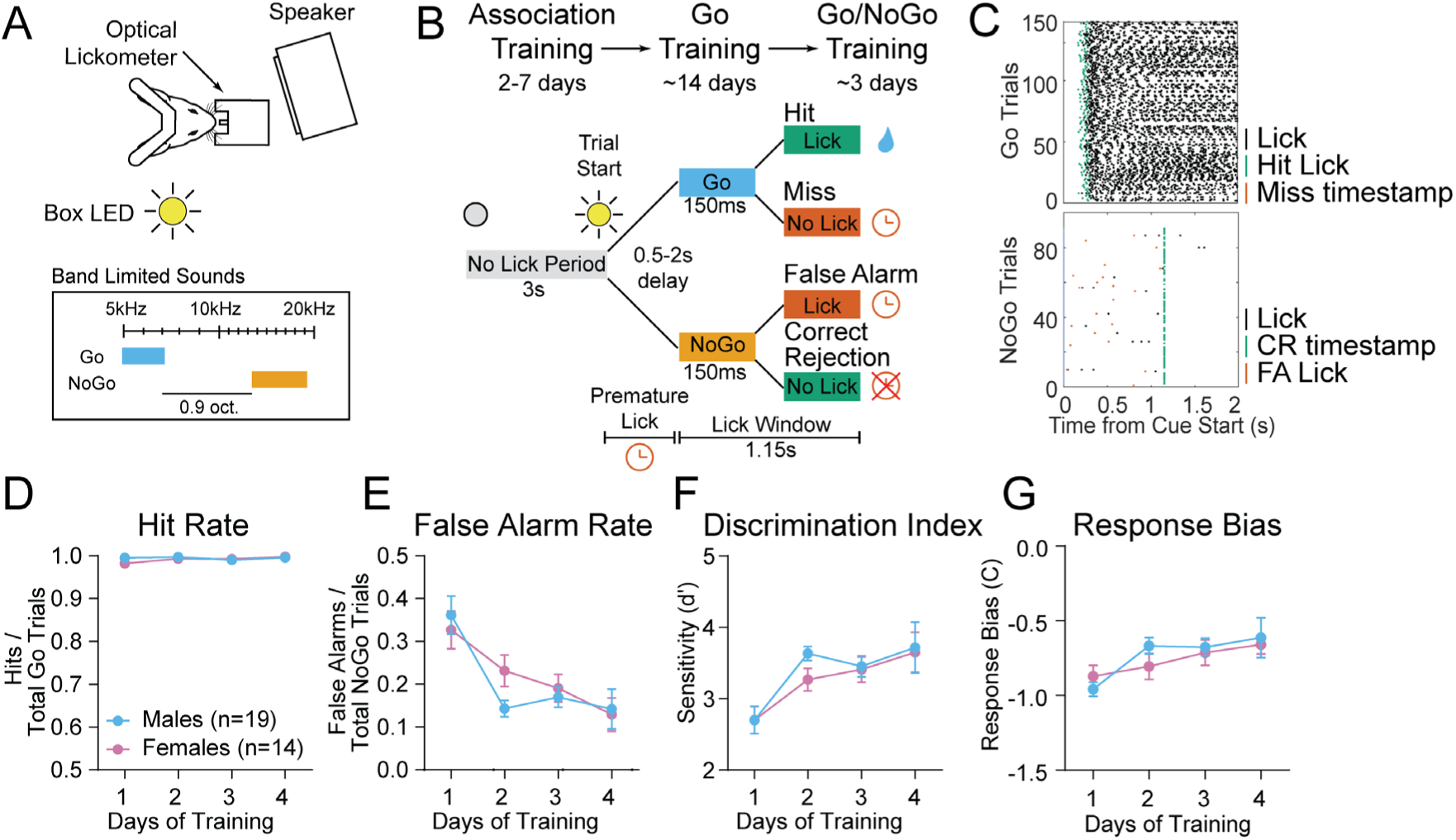
An auditory Go/NoGo task to study sensory-driven inhibitory control. (A) Schematic of behavioral apparatus and bandlimited auditory stimuli used. (B) Training timeline and trial structure for final Go/NoGo behavioral task. (C) Example lick raster plots from expert mice during Go (above) and NoGo (below) trials. (D) Hit rates across the first 4 days of Go/NoGo training for male (n = 19) and female (n = 14) mice. Symbols signify group means ± SEM. All learning data (D-G) were analyzed using a linear mixed effects model with training day (within-subjects) and sex (between-subjects) as fixed effects and subject as a random effect. There were no significant main effects of either training day (F(2.31, 53.91) = 0.66, p = 0.54; Geisser-Greenhouse correction applied) or sex (F(1, 31) = 1.23, p = 0.28) for hit rates. (E) False alarm rates across the first 4 days of Go/NoGo training for male and female mice. Linear mixed effects model: main effect of training day (F(2.17, 50.59) = 14.31, p < 0.0001; Geisser-Greenhouse correction applied); no main effect of sex (F(1, 31) = 0.53, p = 0.47). (F) d′ values across the first 4 days of Go/NoGo training for male and female mice. Linear mixed effects model: main effect of training day (F(2.27, 52.87) = 14.84, p < 0.0001; Geisser-Greenhouse correction applied), but there was no significant effect of sex (F(1, 31) = 0.76, p = 0.39). (G) Response bias values across the first 4 days of Go/NoGo training for male and female mice. Linear mixed effects model: main effect of training day, F(2.75, 64.04) = 5.92, p = 0.0017; no main effect of sex, F(1, 31) = 0.28, p = 0.60.

Mice were initially trained in trials containing only the rewarded Go sound, with the final stage of training introducing a random subset of non-rewarded NoGo sound trials requiring sound-related inhibitory control (Fig. 1B). Most animals could achieve high discriminative performance (d’ > 2) within 4 days of training, driven by a stable hit rate and declining proportion of false alarms (Fig. 1D-F). While animals generally exhibited a bias towards licking, as indicated by negative response biases, these values became less negative with continued training (Fig. 1G). As no significant sex differences were observed in learning or performance of our Go/NoGo behavior, data from both sexes were pooled together for all subsequent experiments (Fig. 1D-G; Fig. S1).

### SPNs are differentially recruited by task-associated auditory stimuli

To observe SPN subtype recruitment as mice performed our auditory-guided task, we utilized fiber photometry to visualize direct and indirect pathway population activity. We selectively expressed the fluorescent calcium sensor, GCAMP8m, in either D1– or D2-dopamine receptor positive SPNs (dSPNs, iSPNs) by injecting a Cre-dependent AAV construct (AAV-hSyn::FLEX-jGCaMP8m) into the TS of D1Cre or A2ACre mice (Fig. 2A,B; Fig.S2A). To get an initial overview of neuronal activity during expert performance, we aligned photometry traces to sound onset and sorted them based on trial type. We found that initial exposure to high frequency sounds consistently produced larger SPN responses in both subtypes, regardless of whether they were target or non-target stimuli (Fig. S2B,C). Based on our reproducible optic cannula placements (Fig. S2A) and the previously reported tonotopic organization of the TS^58^, this suggested our recordings were largely centered on a high frequency preferring region of TS. Given this, we initiated our photometry analysis on the cohort whose Go sound cue was within the high-frequency domain.

**Figure 2.**
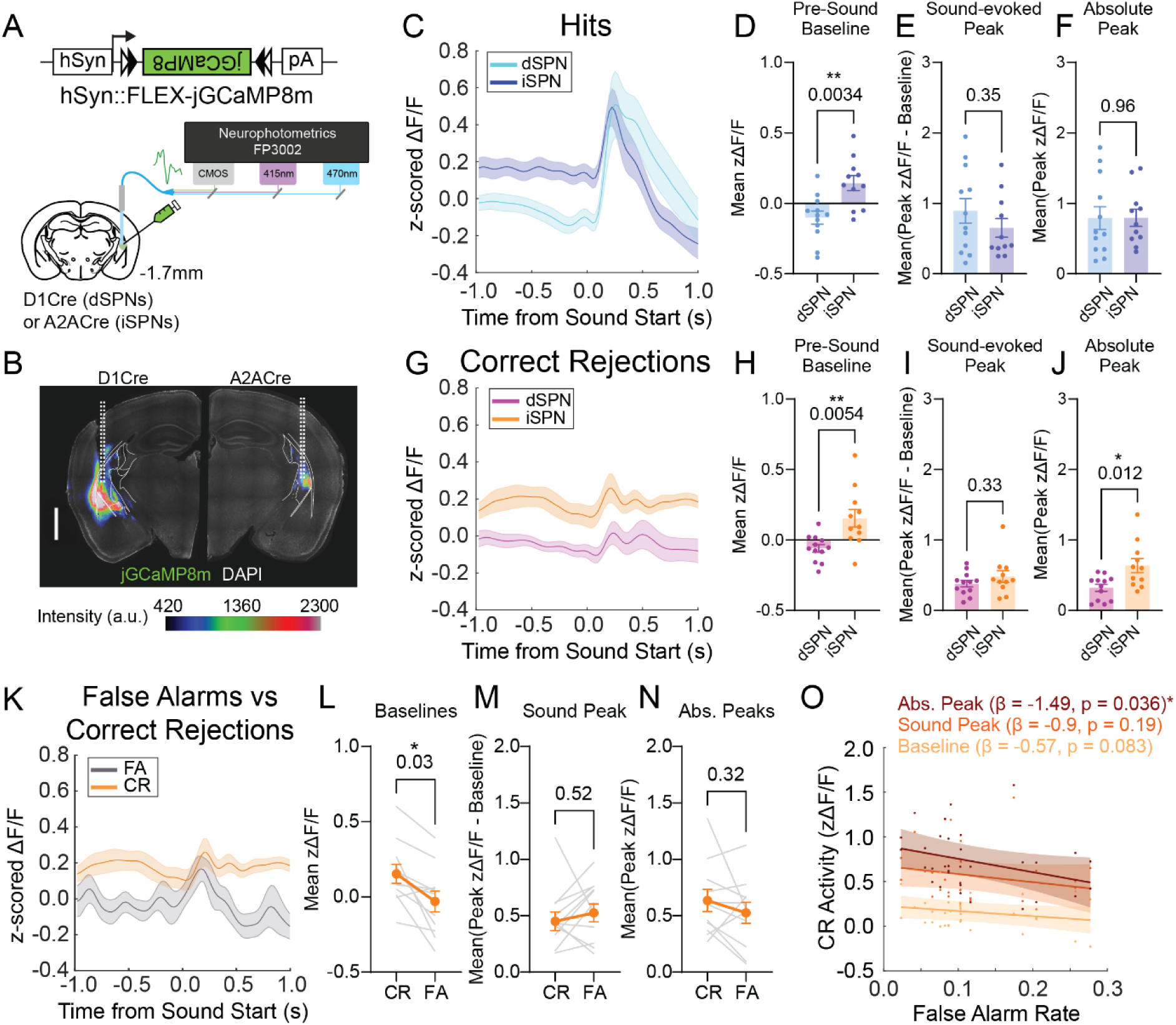
Enhanced recruitment of iSPNs during response inhibition in NoGo trials. (A) Experimental schematic for cell-type specific photometry recordings. (B) Example histological images showing dSPN (left) and iSPN (right) jGCaMP8m fluorescence intensity. Scale bar is 1mm. (C) Group average fluorescence traces aligned to Go sound onset from dSPNs (light blue, n = 6 mice; 12 recording sites) and iSPNs (dark blue, n = 6 mice; 11 recording sites) during hit trials from a single expert session. Shading denotes SEM. (D) Trial-averaged pre-sound baseline fluorescence values for each recording site from expert hit trials. Two-way mixed-effects model (REML) with Pathway and Hemisphere as fixed effects and Subject as a random effect; main effect of Pathway, p = 0.0034**; Hemisphere, p = 0.83; interaction, p = 0.58. (E) Trial-averaged baseline-subtracted peak fluorescence values for each recording site from expert hit trials. Two-way mixed-effects model (REML) with Pathway and Hemisphere as fixed effects and Subject as a random effect; Pathway, p = 0.35; Hemisphere, p = 0.73; interaction, p = 0.54. (F) Trial-averaged absolute peak fluorescence values for each recording site from expert hit trials. Two-way mixed-effects model (REML) with Pathway and Hemisphere as fixed effects and Subject as a random effect; Pathway, p = 0.96; Hemisphere, p = 0.77; interaction, p = 0.40. (G) Group average fluorescence traces aligned to NoGo sound onset from dSPNs (purple, n = 6 mice; 12 recording sites) and iSPNs (orange, n = 6 mice; 11 recording sites) during correct rejection trials from a single expert session. Shading denotes SEM. (H) Trial-averaged pre-sound baseline fluorescence values for each recording site from expert correct rejection trials. Two-way mixed-effects model (REML) with Pathway and Hemisphere as fixed effects and Subject as a random effect; main effect of Pathway, p = 0.0054**; Hemisphere, p = 0.59; interaction, p = 0.54. (I) Trial-averaged baseline-subtracted peak fluorescence values for each recording site from expert correct rejection trials. Two-way mixed-effects model (REML) with Pathway and Hemisphere as fixed effects and Subject as a random effect; Pathway, p = 0.33; Hemisphere, p = 0.90; interaction, p = 0.20.(J) Trial-averaged absolute peak fluorescence values for each recording site from expert correct rejection trials. Two-way mixed-effects model (REML) with Pathway and Hemisphere as fixed effects and Subject as a random effect; main effect of Pathway, p = 0.012*; Hemisphere, p = 0.81; interaction, p = 0.47. (K) Group average fluorescence traces aligned to NoGo sound onset from iSPNs (n = 6 mice; 11 recording sites) during correct rejection (orange) and false alarm (gray) trials in a single session. Shading denotes SEM. (L) Trial-averaged pre-sound baseline fluorescence values for each recording site from expert NoGo trials. Two-way mixed-effects model (REML) with Trial Type and Hemisphere as fixed effects and Subject as a random effect; main effect of Trial Type, p = 0.0298*; Hemisphere, p = 0.88; interaction, p = 0.13. (M) Trial-averaged baseline-subtracted peak fluorescence values for each recording site from expert NoGo trials. Two-way mixed-effects model (REML) with Trial Type and Hemisphere as fixed effects and Subject as a random effect; Trial Type, p = 0.52; Hemisphere, p = 0.81; interaction, p = 0.21. (N) Trial-averaged absolute peak fluorescence values for each recording site from expert NoGo trials. Two-way mixed-effects model (REML) with Trial Type and Hemisphere as fixed effects and Subject as a random effect; Trial Type, p = 0.32; Hemisphere, p = 0.81; interaction, p = 0.47.(O) Relationships between correct rejection neural activity metrics (z-scored ΔF/F) and false alarm (FA) rate across sessions. Lines show linear mixed-effects fits with shaded 95% CIs. Points are individual recording sites per session (6 mice, 11 sites, 3 sessions/site; n = 33 observations per model). Models fit by maximum likelihood: Activity ∼ FA Rate + (1 | Rec Site). Slope estimates (β): Absolute peak –1.49 (p = 0.036)*, Sound peak –0.90 (p = 0.19), Baseline –0.57 (p = 0.083).

We observed sound-evoked activity in the direct and indirect pathway populations for both correct target responses (hits; Fig. 2C-F) and correct non-target rejections (correct rejection, CR; Fig. 2G-J). We quantified three components of the photometry signal including the pre-sound baseline, the sound-evoked peak (i.e. baseline subtracted or peak prominence) and the absolute population peak (a combination of baseline and sound-evoked activity). For hit trials, the absolute peak activity after Go sound presentation did not change over the course of training (Fig. S2F,G). Additionally, both striatal pathways showed robust activation in hit trials, with similar peak responses for dSPN and iSPN subtypes (Fig. 2E,F). This may indicate an organized involvement of both striatal pathways for initiating responses to reward-related target stimuli.

During correct rejection trials, which engage inhibitory control mechanisms to the stimulus, sound related activity in both pathways was reduced compared to hit trials (Fig. S2D,E). While this is likely due to the underlying striatal tonotopy (with low-frequency NoGo sounds evoking smaller responses in a high-frequency predominant area), we noted that initial exposure to the NoGo stimulus elicited greater responses that diminished over the course of training (Fig. S2H,I). In expert mice, unlike the concurrent SPN activity observed during hit trials, the absolute peak following NoGo sound onset was significantly greater in iSPNs than in dSPNs during correct rejection trials (Fig. 2I,J). We also noted that the differences between hit and CR trial peaks were smaller for iSPNs (compare S2D,E). Furthermore, we observed SPN-subtype specific differences in baseline activity leading up to sound onset, with iSPNs having significantly higher baseline activity on both hit and CR trials (Fig 2D,H). While we observed similar trends in mice trained on the low-frequency Go stimulus, differences between CR activity in SPN subtypes did not reach statistical significance (Fig. S3).

To further investigate potential iSPN contributions to inhibitory control, we compared the photometry signals for mice that correctly withheld (CR) or incorrectly responded (false alarm, FA) to the non-target tone. While iSPNs exhibited a significant reduction in baseline activity during false alarm trials compared to correct rejections, the sound-evoked and absolute peak activity following the NoGo sound did not significantly change between false alarm trials and correct rejection trials (Fig 2K-N). Given the possible contributions of baseline activity shifts in iSPNs as a mechanism of sound-driven inhibitory control, we examined this phenomenon further. We first aligned our traces to the start of every trial, finding that the enhanced baseline activity appeared at house light onset, which marked the trial start (Fig.S2J). As animals are required to withhold licking for 3s prior to the trial start, this suggests that the baseline iSPN activity is not directly related to a general withholding of licking. Furthermore, we found that this elevated baseline emerged during the earliest stage of training, before premature licking was penalized or licking behavior during the trial influenced reward availability (Fig. S2K, L). This suggests that while this elevated iSPN baseline activity may contribute to inhibitory control in some regard, it is not necessarily exclusive to it.

We therefore sought to investigate whether sound-related iSPN activity could best predict inhibitory control performance, as the decreased iSPN baseline activity—though correlated with false alarm trials—did not appear to be specific to inhibitory control. To do so, we correlated our iSPN activity measures during CR trials with inhibitory control performance across multiple sessions, as measured by false alarm rate (Fig. 2O). Although baseline iSPN activity showed a negative trend with false alarm errors, this relationship did not reach statistical significance, further suggesting that baseline iSPN activity alone may not directly mediate inhibitory control of sound-driven behavior. Similarly, the sound-evoked response during CR trials did not correlate with NoGo performance. In contrast, only the absolute peak activity—which encompasses both baseline and sound-evoked components—demonstrated a significant negative correlation with false alarm error rates. This supports the idea that absolute peak iSPN activity recruited during correct NoGo trials can predict overall inhibitory control performance for a given session, suggesting the involvement of a sound-related iSPN population that mediates inhibitory control over sound-driven actions.

Taken together, our population imaging shows subtype specific activity modulation which diverges for target and non-target sounds. On Go trials, we find similar absolute peak activity for both cell-types, suggesting that a coordinated effort between pathways may mediate reward-related responses to target sounds. However on NoGo trials, we find greater iSPN than dSPN activity aligned to the onset of the NoGo sound, indicating potential iSPN involvement for inhibitory control to inappropriate or non-target stimuli. Furthermore, we find that while iSPNs exhibit a behaviorally correlated, tonic baseline activity preceding sound presentation, this baseline does not directly mediate inhibitory control. Rather it appears to facilitate it, as baseline activity is reduced in false alarm trials and likely contributes to the absolute peak activity observed after sound presentation. Finally, overall sound-related iSPN activity in correctly rejected NoGo trials seems to predict non-target preformance across sessions. Together these data suggest that iSPN activity both before and during sound presentation seem to differentially support conservative strategic approaches or proactive inhibitory control for auditory driven action.

### Global inhibition of TS SPNs selectively impairs lick responses to target stimuli

To causally probe the contributions of SPN-specific activity patterns to behavioral performance, we first used muscimol to simultaneously inhibit both SPN pathways (Fig.3). We bilaterally injected the GABA_A_ receptor agonist into the TS during expert Go/NoGo sessions, one day after administering PBS vehicle injections as a paired control (Fig. 3A). Muscimol administration significantly and reversibly suppressed responses to the target sound, while false alarm response rate remained unchanged (Fig.3B,C). This impairment in reward-related responding significantly reduced discriminative performance and resulted in a shift in an overall response bias towards withholding (Fig. 3D,E). The effects of muscimol administration on Go/NoGo behavior had largely disappeared 2.5 hours following initial injection (Fig 3B-E).

**Figure 3.**
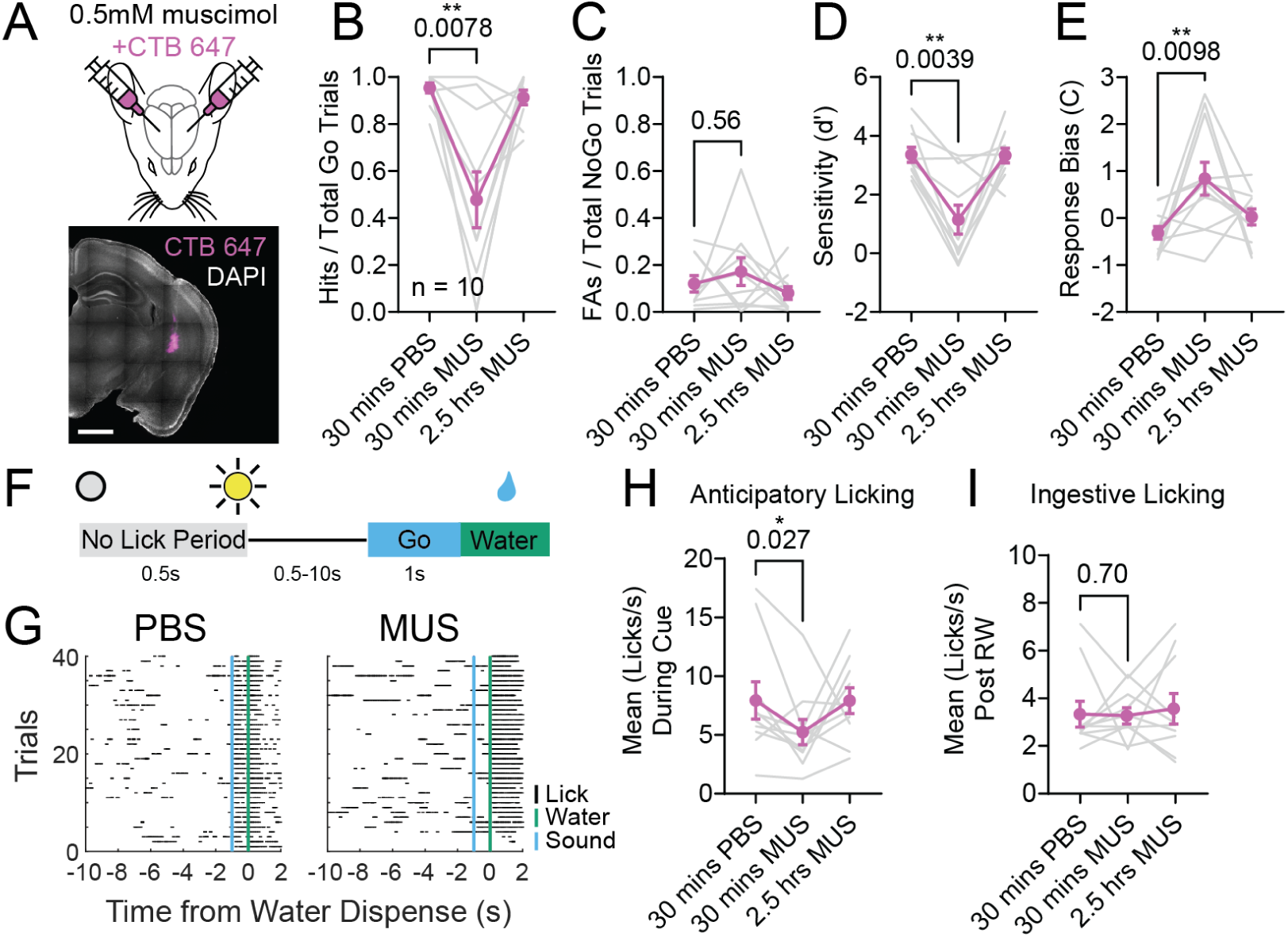
Coordinated activity of both SPN pathways are required for appropriate target sound responses. (A) Experimental schematic for bilateral vehicle/drug administration with example histological image showing target site of muscimol release. Scale bar is 1mm. (B) Hit rate for each animal 30 mins post PBS vehicle administration, 30 mins post muscimol administration and 2.5 hours post muscimol administration. Gray lines connect individual mice across conditions; magenta symbols show mean ± SEM (n = 10). Comparison of 30 min MUS vs 30 min PBS: two-tailed Wilcoxon matched-pairs signed-rank test (exact): W = −43, p = 0.0078**. (C) False alarm rate for each animal 30 mins post PBS vehicle administration, 30 mins post muscimol administration and 2.5 hours post muscimol administration. Gray lines connect individual mice across conditions; magenta symbols show mean ± SEM (n = 10). Comparison of 30 min MUS vs 30 min PBS: two-tailed Wilcoxon matched-pairs signed-rank test (exact): W = 13, p = 0.56. (D) Discrimination index for each animal 30 mins post PBS vehicle administration, 30 mins post muscimol administration and 2.5 hours post muscimol administration. Gray lines connect individual mice across conditions; magenta symbols show mean ± SEM (n = 10). Comparison of 30 min MUS vs 30 min PBS: two-tailed Wilcoxon matched-pairs signed-rank test (exact): W = −53, p = 0.0039**. (E) Response bias for each animal 30 mins post PBS vehicle administration, 30 mins post muscimol administration and 2.5 hours post muscimol administration. Gray lines connect individual mice across conditions; magenta symbols show mean ± SEM (n = 10). Comparison of 30 min MUS vs 30 min PBS: two-tailed Wilcoxon matched-pairs signed-rank test (exact): W = 56, p = 0.0098**. (F) Schematic for the non-operant session that animals ran in immediately prior to each Go/NoGo session. (G) Example lick rasters from a mouse 30 minutes after PBS administration (left) and 30 minutes after muscimol administration (right). Licks are denoted in black, sound start timestamp in blue and water dispense timestamp in green. (H) Average lick rate for each animal during Go sound presentation in the non-operant session for each experimental timepoint. Gray lines connect individual mice across conditions; magenta symbols show mean ± SEM (n = 10). Comparison of 30 min MUS vs 30 min PBS: two-tailed Wilcoxon matched-pairs signed-rank test (exact): W = –43, p = 0.027*. (I) Average lick rate for each animal after water presentation in the non-operant session for each experimental timepoint. Gray lines connect individual mice across conditions; magenta symbols show mean ± SEM (n = 10). Comparison of 30 min MUS vs 30 min PBS: two-tailed Wilcoxon matched-pairs signed-rank test (exact): W = 9, p = 0.70.

To rule out motivational deficits or impairments in overall lick ability, prior to each Go/NoGo session we also ran animals in a non-operant version of the behavior where licking had no influence on reward delivery and instead reward was dispensed following a variable duration after trial start (Fig. 3F). This random reward delivery was consistently preceded by the presentation of the Go sound. During PBS sessions, animals predictably licked during the Go stimulus in anticipation of reward (Fig. 3G, H). While muscimol administration significantly reduced this anticipatory licking, the ingestive lick rate after reward dispensing was not significantly altered (Fig. 3G-I). This indicates that although animals exhibited an intact motivational drive for water reward and overall ability to lick to ingest reward, licking in response to the target stimulus was specifically disrupted. Taken together with our photometry observations of concurrent SPN recruitment for hit trials, this suggests that coordinated activity between both SPN subtypes within the TS is required for proper reward-related responses to auditory stimuli.

### TS dSPN manipulations alone minimally impact sound-driven behavior

To examine the temporal contributions of SPN subtypes to auditory driven action control, we utilized cell-type specific optogenetic inhibition and excitation (Fig. S4). Given our prior data, we wondered whether inhibition of dSPNs during target sounds would decrease hit trials and whether inhibition during non-target sounds would decrease false alarms. We gained inhibitory optogenetic control of dSPNs by crossing D1Cre and Ai40D mice, which allow Cre-dependent expression of the inhibitory opsin ArchT3.0 (Fig. S4A). The resulting fluorescence patterns was consistent with previously reported dopamine D1 receptor expression patterns and demonstrated robust substantia nigra fluorescence from axonal projections (Fig.S4A). D1-Cre; R26R^AI40/+^ mice and their littermate controls (R26R^AI40/+^) were bilaterally implanted with fiber optic cannulae in TS and SPNs were inhibited during sound presentation on a randomized subset (25%) of Go/NoGo trials using 250ms. of 10mW, 532 nm continuous green light stimulation (Fig. S4B, K; Fig. S6A). Surprisingly, we found that dSPN inhibition specifically at target sound presentation did not alter the proportion of hit trials nor shift the distribution of hit reaction times (Fig.S4C,E,L). Furthermore, inhibition of dSPNs during non-target sounds did not decrease the FA rate, although a floor effect could be possible (Fig.S4C). We also observed a modest increase in FA reaction times and small decrease in response bias (Fig.S4D,E,L).

We next tested whether dSPN excitation was sufficient to alter behavior (Fig.S4F-J,M). We expressed the red-shifted excitatory opsin, ChrimsonR in dSPNs via injection of a Cre-dependent AAV construct (AAV8-hSyn-FLEX-ChrimsonR-tdTom) with bilateral fiber optic cannulae implants (Fig.S4F). We selectively activated dSPNs during sound presentation of randomly selected Go/NoGo trials (25%) using 5mW, 625nm red, 20Hz pulsed light (Fig.S4G). The high hit rate in our task precluded strong conclusions about the ability of dSPN to increase responding to target tones, but excitation did produce a small but reliable reduction in hit reaction times (Fig.S4H,J,M). Surprisingly, the addition of dSPN activity during non-target tones did not increase false alarms or reaction times (Fig.S4H,J,M). Moreover, overall d’ and response bias did not significantly change during D1-SPN activation (Fig.S4I). Together, these modest behavioral effects for dSPN perturbations may suggest the involvement of coordinated SPN subtype activity in reward-related responding, interference from concurrent lateral inhibition of iSPNs, or redundancy in the locus of behavioral control for reward-related responding^26,59^.

### Perturbations of TS iSPN activity alters sound-driven action control

Our photometry experiments suggested that iSPNs in this striatal region may support auditory driven response control, with recruitment of iSPNs potentially suppressing inappropriate licking in response to the NoGo stimulus. To test this, we first decided to inhibit iSPN activity during sound presentation, hypothesizing that this would increase the proportion of false alarm responding (Fig.4A-E). We crossed A2A-Cre and R26R^Ai40D^ mice and observed robust ArchT3.0 expression of TS iSPNs in addition to some non-neuronal labeling in cortex and hippocampus (Fig.4A, Fig.S5A). When we randomly inhibited iSPNs during non-target sound presentation we found a robust increase in false alarm rate which was matched with a decrease in reaction time to the NoGo stimulus (Fig.4C, E, Fig.S5B). While inhibition of iSPNs during target sounds did not change the hit rate, we did observe a consistent reduction in the reaction time associated with target responding (Fig.4E, Fig.S4B). Overall, impaired NoGo performance was associated with a significant reduction in discriminative sensitivity, accompanied by an increased bias toward licking responses (Fig. 4D). This pattern suggests a shift toward a more impulsive response strategy, in which animals were less likely to withhold licking in the presence of the NoGo stimulus. No light related effects were observed in Cre-negative controls (Fig. S6A-D, H). To further probe the temporal constraints of iSPN function, we performed another 250ms iSPN inhibition after the sound presentation, timed to the average onset of the lick response (Fig.S5C). We found that this temporally delayed manipulation had consistently smaller impacts on the production of false alarms and accompanying signal detection metrics, while having no impact on response times for either trial type (Fig.S5D-G). Once again, there were no robust light-related effects observed in Cre-negative controls during post-sound sessions (Fig. S6E-G, I).

**Figure 4.**
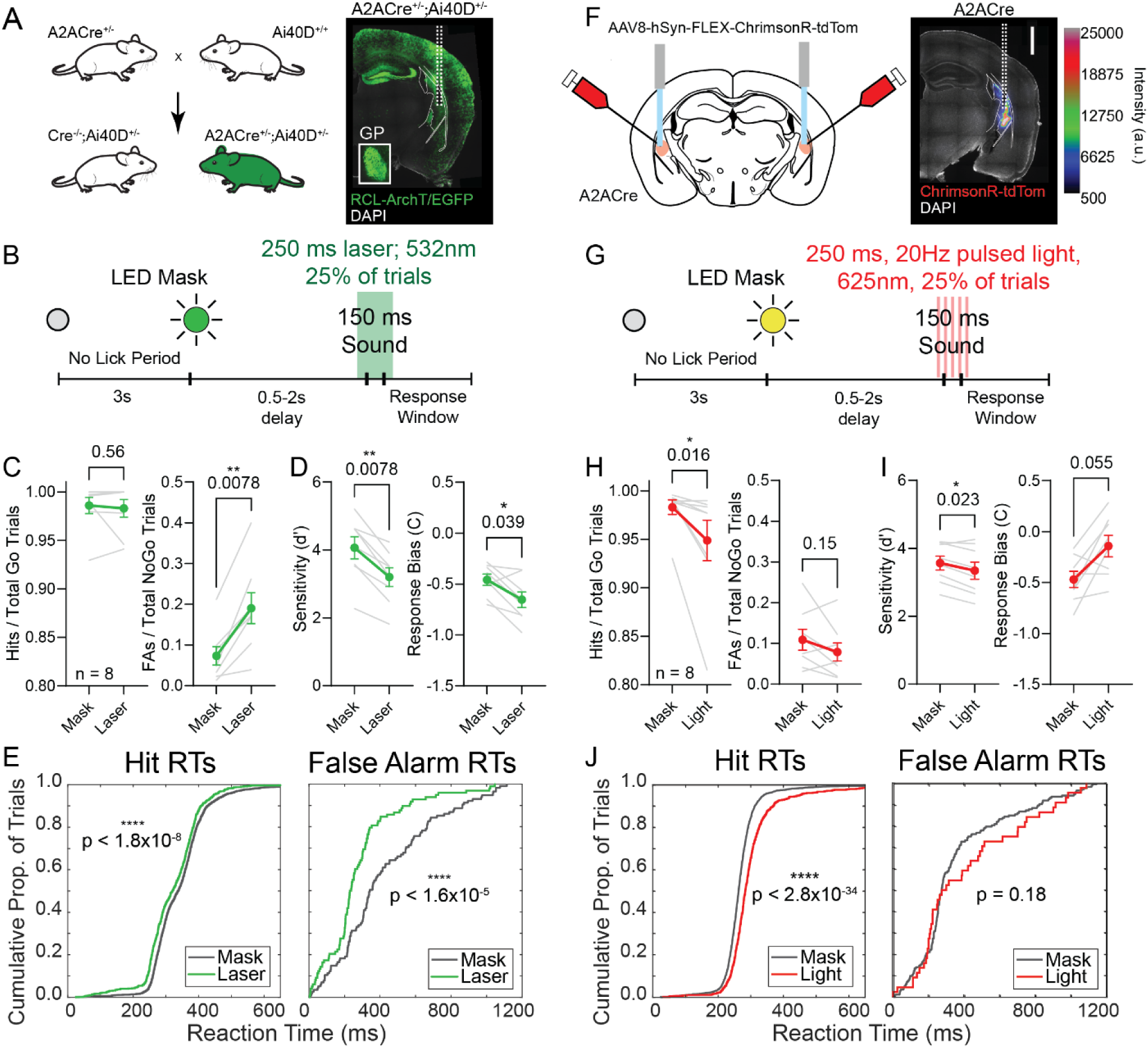
The indirect pathway is involved in mediating correct rejection of the non-target stimulus. (A) Experimental breeding strategy to generate transgenic mice expressing ArchT in iSPNs with example histology image. Inset shows iSPN terminal expression in globus pallidus. (B) Schematic showing time course for laser-mediated inhibition of iSPNs during sound presentation. (C) Hit (left) and false alarm (right) rates for mask control versus laser mediated inhibition trials. Gray lines connect individual mice across conditions; green symbols show mean ± SEM (n = 8). Comparison of mask vs laser hit rates: two-tailed Wilcoxon matched-pairs signed-rank test (exact): W = 7, p = 0.56. Comparison of mask vs laser false alarm rates: two-tailed Wilcoxon matched-pairs signed-rank test (exact): W = 36, p = 0.0078**. (D) Discrimination sensitivity (left) and response bias (right) values for mask control versus laser mediated inhibition trials. Gray lines connect individual mice across conditions; green symbols show mean ± SEM (n = 8). Comparison of mask vs laser d’ values: two-tailed Wilcoxon matched-pairs signed-rank test (exact): W = –36, p = 0.0078**. Comparison of mask vs laser response biases: two-tailed Wilcoxon matched-pairs signed-rank test (exact): W = –30, p = 0.039*. (E) Cumulative distributions for hit (left) and false alarm (right) reaction times (RT) during mask and laser inhibition conditions. Curves pool trials from multiple sessions across 8 mice [Mask: n_hit_ = 2447, n_FA_ = 115; Laser: n_hit_ = 720, n_FA_ = 98]. Differences between distributions were tested with two-sample Kolmogorov-Smirnov tests (two-sided): Hits, D = 0.13, p = 1.8*10^−8^; False Alarms, D = 0.33, p = 1.6*10^−5^. (F) Experimental schematic to express the excitatory, red-shifted opsin ChrimsonR in iSPNs with example histology image. Scale bar is 1mm. (G) Schematic showing time course for light-mediated excitation of iSPNs during sound presentation. (H) Hit and false alarm rates for mask control versus light mediated excitation trials. Gray lines connect individual mice across conditions; red symbols show mean ± SEM (n = 8). Comparison of mask vs light hit rates: two-tailed Wilcoxon matched-pairs signed-rank test (exact): W = –34, p = 0.016*. Comparison of mask vs light false alarm rates: two-tailed Wilcoxon matched-pairs signed-rank test (exact): W = –22, p = 0.15. (I) Discrimination sensitivity and response bias values for mask control versus light mediated excitation trials. Gray lines connect individual mice across conditions; red… …symbols show mean ± SEM (n = 8). Comparison of mask vs light d’ values: two-tailed Wilcoxon matched-pairs signed-rank test (exact): W = –32, p = 0.023*. Comparison of mask vs light response biases: two-tailed Wilcoxon matched-pairs signed-rank test (exact): W = 28, p = 0.055. (J) Cumulative distributions for hit and false alarm reaction times (RT) during mask and light excitation conditions. Curves pool trials from multiple sessions across 8 mice [Mask: n_hit_ = 2525, n_FA_ = 831; Light: n_hit_ = 172, n_FA_ = 44]. Differences between distributions were tested with two-sample Kolmogorov-Smirnov tests (two-sided): Hits, D = 0.25, p = 2.8*10^-34^; False Alarms, D = 0.18, p = 0.18.

Finally, we tested the impact of ectopically activating iSPNs by using bilateral injections of AAV8-hSyn-FLEX-ChrimsonR-tdTom into A2a-Cre mice (Fig. 4F, G). We hypothesized that augmenting iSPN activity could decrease FA and target sound responding, promoting a more conservative response bias. We found that excitation of iSPNs during Go trials significantly decreased the hit responses and increased reaction times (Fig. 4H, J). We did not observe a significant change in false alarm rate or false alarm reaction time with iSPN activation during the NoGo stimulus. This selective impairment in the hit rate resulted in a small but consistent decrease in sensitivity and a trending increase in response bias consistent with a reduced bias toward licking (Fig. 4I). Taken together, these data suggest that iSPN activity mediates response withholding for non-target sounds and can support more conservative response strategies to sounds.

### Optogenetic manipulation of TS iSPNs biases licking after sound-action association forms

To examine whether our iSPN manipulations are modulating licking as a learned behavioral response or directly triggering motor output, we manipulated iSPN activity outside the direct context of our behavioral paradigm (Fig. S7). We examined two training timepoints--the first occurring before animals had ever undergone any auditory training (Naïve Opto), and the second session occurred after animals had learned to successfully restrict their lick responses to the Go sound (Go Trained Opto) (Fig. S7A,B). In these sessions, auditory stimuli were omitted entirely, and licking did not influence reward delivery. To encourage free licking, there were blocks of trials during which water was randomly dispensed (Fig. S7B). These blocks were then followed by stimulation blocks where 1s of light stimulation randomly occurred. Licking during these blocks were recorded and analyzed to determine how SPN stimulation impacts free licking probability.

When animals were naive to any type of sound training, iSPN stimulation did not alter free licking rate (Fig. S7C,D,G,H). However, after animals had learned to restrict their licking to be used as a response to the Go stimulus, iSPN manipulation demonstrated opponent effects on free licking rate. iSPN excitation significantly decreased the lick rate, whereas iSPN inhibition significantly increased the lick rate (Fig. S7E,I). Notably, these changes in lick rate were not consistently observed across trials, as even animals displaying the strongest phenotype did not show reliable alterations in licking patterns indicative of direct motor program execution (Fig. S7F,J). Therefore, it seems that indirect pathway neurons can bidirectionally bias free licking probability even in the absence of sound. However, this only occurs after animals have been trained to use the act of licking as a sound-driven behavioral response.

### Neurexin1α-associated disruptions of corticostriatal input to TS iSPNs is related to impaired inhibitory control

Thus far, our circuit analyses suggest that impairments of iSPN function within the TS could disrupt the ability to inhibit responses to non-target stimuli. We next sought to investigate whether synaptic dysfunction within this circuit could contribute to the inhibitory control impairments frequently seen in NDDs, using a genetic model with previously established iSPN corticostriatal alterations. Neurexin1α (Nrxn1α) is a synaptic adhesion molecule whose disruption is associated with a range of neurodevelopmental and psychiatric disorders including autism, Tourette syndrome, schizophrenia, and attention-deficit hyperactivity disorder^45,49–56,60,61^. Prior acute slice studies have shown that dorsomedial prefrontal cortical neurons in both Nrxn1α^+/−^ and Nrxn1α ^-/-^ mice exhibit a reduction in excitatory synaptic strength for connections onto iSPNs, but not dSPNs of the dorsomedial striatum (DMS)^45^.

We first investigated whether this synaptic deficit could also be observed in the TS. To quantitatively measure cortically derived synaptic strength to the tail, we aimed to express the excitatory opsin ChR within all excitatory cortical inputs in Nrxn1α mutant mice. To accomplish this, we took advantage of the previously described haploinsufficiency of Nrxn1α (Nrxn1α^+/−^ and Nrxn1α ^-/-^ exhibit similar magnitude reductions in synaptic strength in DMS)^45^, crossing Nrxn1α^+/−^ mice to Nex^Cre/Cre^; R26R^AI32/AI32^; D1-Tomato mice to generate either Nrxn1α^+/+^ or Nrxn1α^+/−^ mice with a single copy of both Nex^Cre/+^ and R26R^AI32/+^, a Cre-sensitive ChR134-EYFP reporter line (Fig.5A). This allows us to systematically compare the strength of corticostriatal synaptic inputs from neocortex between Nrxn1α genotypes, by leveraging a genetics-based normalization of channel-rhodopsin expression^62^. Using fluorescent guided patching of the D1-Tom allele, we recorded from putative iSPNs identified by the absence of tdTOM.

**Figure 5.**
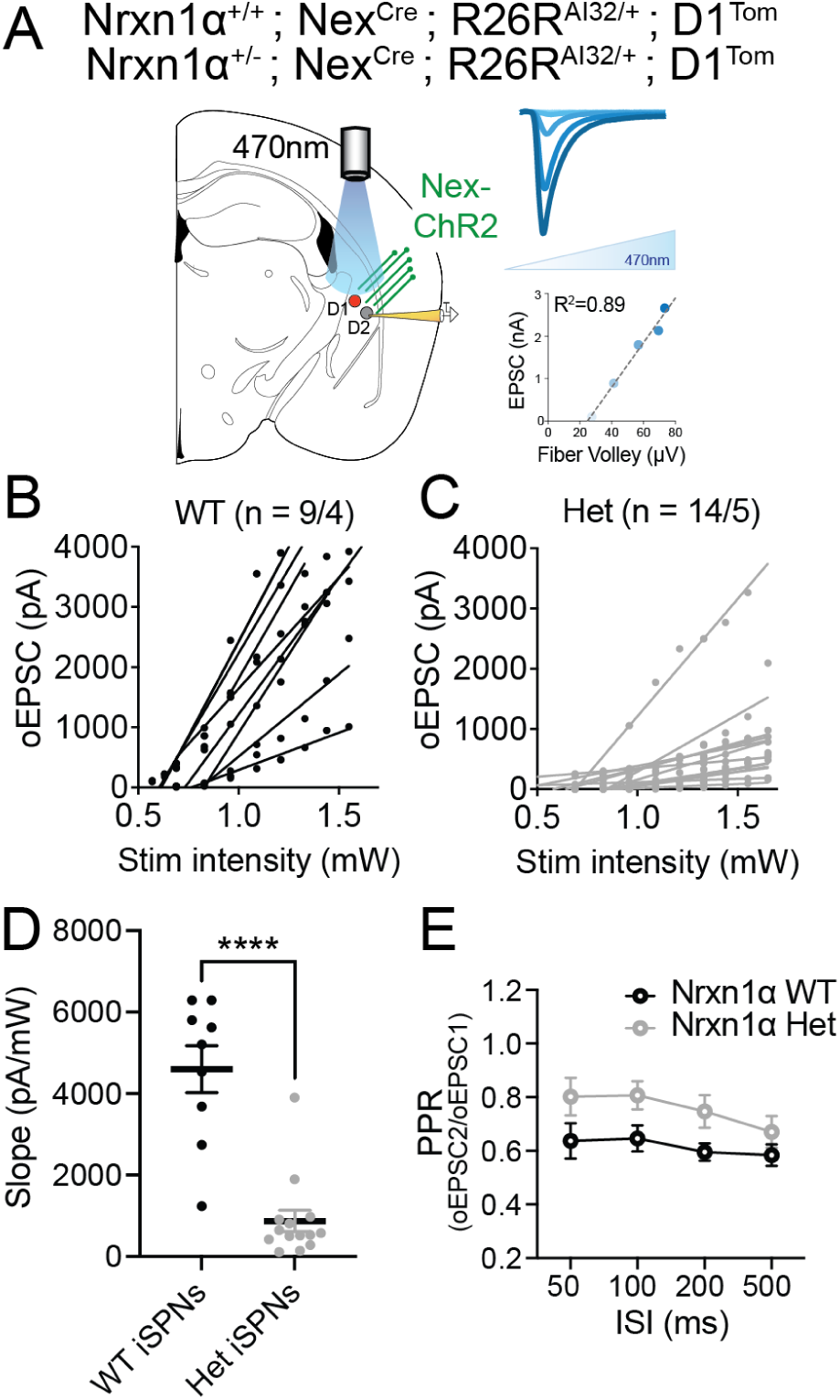
Nrxn1α mutant mice exhibit impaired cortical recruitment of iSPNs within the TS. (A) Experimental strategy to generate Nrxn1α transgenic mice expressing tdTomato in dSPNs as well as ChR2 in cortical inputs. Patched iSPNs were identified as tdTomato negative SPNs. (Right, top) Inward excitatory synaptic currents increased with greater light intensity and synaptic strength could be approximated from a linear fit (right, bottom). (B) Optically evoked EPSCs linear fits for iSPNs from Nrxn1α wildtype mice (n = 9 cells/4 mice). (C) Optically evoked EPSCs linear fits for iSPNs from Nrxn1α heterozygous mice (n = 14 cells/5 mice). (D) Comparison of the slopes from the fits shown in B and C. Individual slopes represented by dots with group means ± SEM shown. Welch’s two-sample t-test (two-sided): WT (n = 9, xō = 4600) vs Het (n = 14, xō = 873), t(11.3) = 5.87, p < 0.0001****; Δ(Het−WT) = −3728 ± 636 SEM; 95% CI [−5122, −2333]. (E) Paired pulse ratios for optical stimulation of iSPNs from Nrxn1α wildtype versus Nrxn1α heterozygous mice across increasing inter stimulus intervals (ISI). Two-way ANOVA (ISI, Genotype). Genotype effect: F(1,84) = 11.82, p = 0.0009; no ISI effect: F(3,84) = 1.29, p = 0.28; no interaction: F(3,84) = 0.20, p = 0.896.

We recorded pharmacologically isolated excitatory synaptic currents in whole cell voltage clamp configuration, subjected each slice to increasing light intensity to sequentially recruit a larger proportion of cortical inputs (Fig.5A). We quantified this input-output function by fitting evoked currents to a linear model and reporting slope as a measure of synaptic strength. Consistent with prior work in DMS, we found that iSPNs in the TS had a dramatically reduced synaptic strength slope in Nrxn1α^+/−^ mice as compared to Nrxn1α^+/+^ (Fig.5B-D). Furthermore, we found that the reduction in TS iSPN synaptic drive was accompanied by a broad increase in paired-pulse ratio, suggestive of a decrease in presynaptic release probability (Fig. 5E). These findings extend prior observations in the DMS to the striatal tail, demonstrating that Nrxn1α haploinsufficiency results in a robust reduction in cortical excitatory drive onto iSPNs throughout the dorsal striatum.

Given the evidence that reductions in iSPN activity could increase inappropriate responding to non-target sounds (Fig.4), together with observations that cortical synaptic transmission to TS iSPNs is impaired (Fig.5), we hypothesized that mice with mutations in Nrxn1α would show deficits in sound-driven response inhibition. To test this, we trained Nrxn1α^+/+^ (WT) mice and littermate Nrxn1α^-/-^ (KOs) mice in our Go/NoGo task to investigate any deficits in inhibitory control to the NoGo stimulus. Here we elected to use the sensitized genetic background of Nrxn1α KOs rather than the more clinically relevant Nrxn1α^+/−^ model as prior mouse work has shown variable penetrance of numerous behavioral phenotypes for Nrxn1α^63^. During Go/NoGo training, although KO mice demonstrated similar hit rates to their littermate controls, KOs exhibited greater false alarm rates throughout initial training (Fig.6A,B). This corresponded with a decrease in the discriminative performance for KOs as well as greater bias towards responding compared to WTs, regardless of the stimuli (Fig.6C,D).

**Figure 6.**
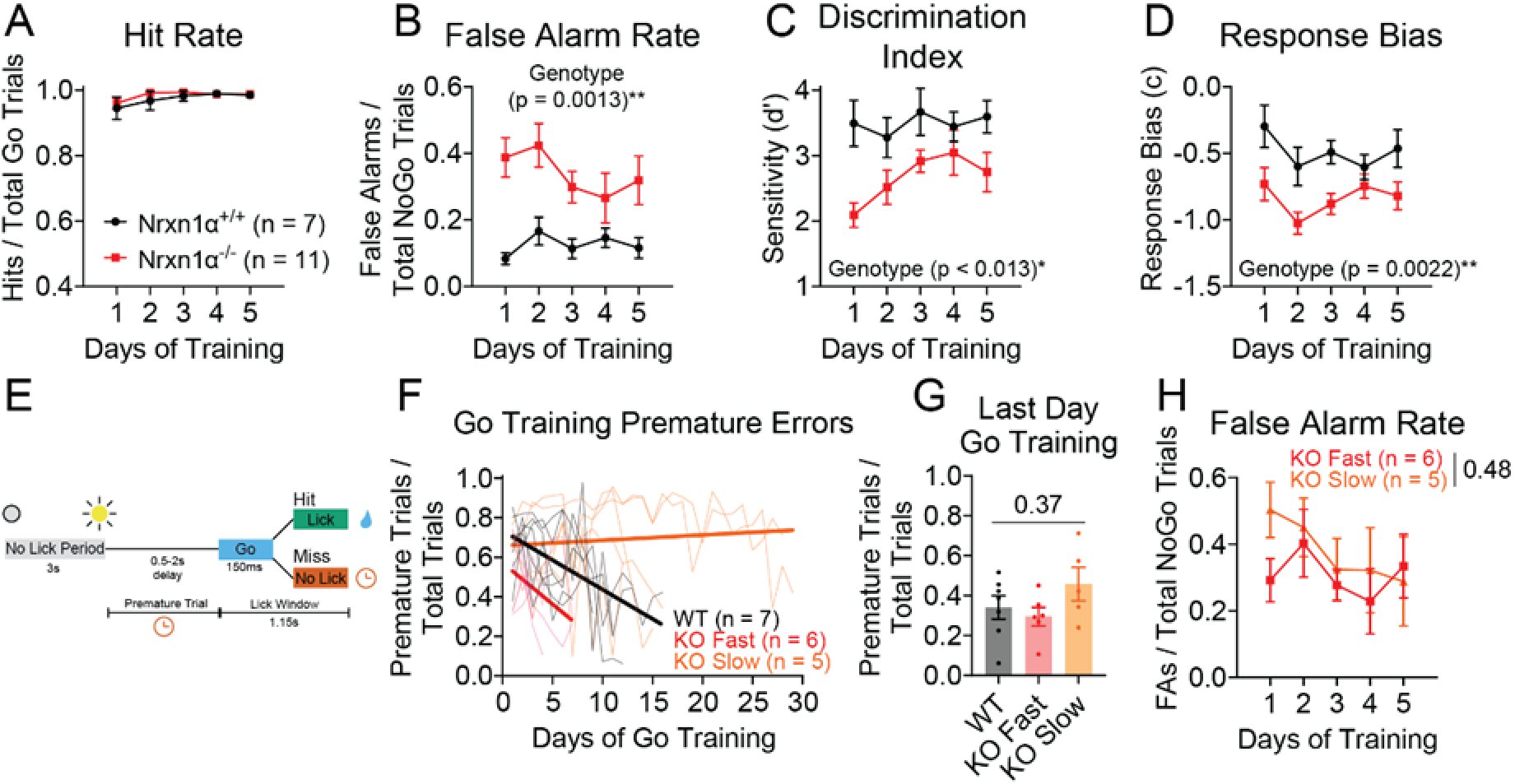
Nrxn1α homozygous knockout mice exhibit specific deficits in withholding responses to non-target sounds. (A) Hit rates for Nrxn1α wildtype (n = 7) and Nrxn1α knockout (n = 11) mice across Go/NoGo training. Symbols denote mean ± SEM. Analyzed using a two-way mixed-effects model (REML) with fixed effects of Training (day) and Genotype, and a random intercept for Subject; sphericity not assumed (Greenhouse–Geisser ε = 0.4989). Main effect of Training: F(1.996, 30.44) = 3.49, p = 0.043*; no Genotype effect: F(1,16) = 0.47, p = 0.50; no interaction: F(4,61) = 0.36, p = 0.84. (B) False alarm rates for Nrxn1α wildtype and Nrxn1α knockout mice across Go/NoGo training. Symbols denote mean ± SEM. Analyzed using a two-way mixed-effects model (REML) with fixed effects of Training (day) and Genotype, and a random intercept for Subject; sphericity not assumed (Greenhouse–Geisser ε = 0.75). Training: F(3.02, 45.98) = 1.21, p = 0.32; Genotype effect: F(1, 16) = 15.25, p = 0.0013**; interaction: F (4, 61) = 1.07, p = 0.38. (C) Discriminative sensitivity for Nrxn1α wildtype and Nrxn1α knockout mice across Go/NoGo training. Symbols denote mean ± SEM. Analyzed using a two-way mixed-effects model (REML) with fixed effects of Training (day) and Genotype, and a random intercept for Subject; sphericity not assumed (Greenhouse–Geisser ε = 0.82). Training: F(3.26, 49.77) = 2.37, p = 0.077; Genotype effect: F(1, 16) = 7.898, p = 0.013*; interaction: F(4, 61) = 1.595, p = 0.19. (D) Response bias for Nrxn1α wildtype and Nrxn1α knockout mice across Go/NoGo training. Symbols denote mean ± SEM. Analyzed using a two-way mixed-effects model (REML) with fixed effects of Training (day) and Genotype, and a random intercept for Subject; sphericity not assumed (Greenhouse–Geisser ε = 0.79). Training: F(3.17, 48.26) = 2.369, p = 0.079; Genotype effect: F(1, 16) = 13.22, p = 0.201C**; interaction: F(4, 61) = 0.69, p = 0.60. (E) Trial schematic for Go training phase. (F) Go training learning curves for Nrxn1α wildtype, Nrxn1α knockout (fast Go learners) and Nrxn1α knockout (slow Go learners). (G) Premature error rates on the last day of Go training for Nrxn1α wildtype (n = 7), Nrxn1α knockout (fast Go leaners; n = 6) and Nrxn1α knockout (slow Go learners; n = 5). Points are individual mice and bars show mean ± SEM. Group comparison by Kruskal-Wallis test (exact): H = 2.086, p = 0.37. (H) Same data shown in B but with the Nrxn1α knockout cohort split by Go learning group. Differences between the two KO cohorts were analyzed using a two-way mixed-effects model (REML) with fixed effects of Training (day) and Go Learning Rate, and a random intercept for Subject; sphericity not assumed (Greenhouse–Geisser ε = 0.69). Training: F(2.74, 22.60) = 1.62, p = 0.21; Go Learning Rate effect: F(1, 9) = 0.55, p = 0.48; interaction: F(4, 33) = 0.82, p = 0.52.

Although previous reports of hyperactivity in Nrxn1α KOs have been inconclusive^57,64–68^, it remains possible that a general increase in impulsive licking, rather than a sound-specific deficit in response inhibition, is driving the elevated false alarm rate observed in these mice. To examine this further, we investigated premature responses arising during the earlier Go training stage (Fig. 6E). Sound delivery in our task was predicated on a variable 0.5-2s window of no licking following trial start, so we decided to use the number of violations of this rule as a proxy for global impulsivity. To reach criteria during the Go training stage, animals were required to reduce premature responses to achieve a *d′* > 2 (comparison of hit to premature response rate). We therefore first examined whether Nrxn1α KO mice required more training sessions to reach criterion performance compared to WT controls. While some Nrxn1α KO mice required more sessions to reach criterion than their WT littermates, approximately half reached expert performance more quickly (Fig.6F). To capture this variability, we classified KO animals as “fast” or “slow” learners based on whether they reached criterion in fewer than 12 sessions (Fig. 6F). Regardless of learning rate, as a group Nrxn1α KO mice did not demonstrate significant differences in premature error rates upon reaching expert criteria (Fig. 6G). Next, we examined whether these fast and slow Go learners within the Nrxn1α KO group differed in their NoGo response inhibition, as fast Go learners might exhibit performance more comparable to WT controls. When we analyzed the proportion of false alarms for these two groups during the final stage of Go/NoGo training, we found that the KO groups were not statistically different in their learning, showing a similar time course and magnitude of increased false alarm rates (Fig. 6H). Taken together, these data suggest that while a subset of Nrxn1α KO mice exhibit generalized motor impulsivity, the entire KO cohort demonstrate a specific impairment in withholding responses to non-target sounds. Furthermore, this deficit in sound-driven response inhibition likely arises, at least in part, from the corticostriatal synaptic disruptions we observed onto iSPNs within the TS.

## Discussion

In this study, we employed an auditory Go/NoGo task to examine the striatal circuitry supporting inhibitory control for sensory-driven behavior, with a particular focus on how disruptions of these mechanisms may contribute to inhibitory control impairments in a genetic neurodevelopmental disorder model. We found that while SPNs from both the direct and indirect pathways apper to mediate the ability to respond appropriately to reward-related cues, we found iSPNs were selectively recruited during correct NoGo trials, supporting their role in suppressing responses to non-target stimuli. This interpretation was further reinforced by optogenetic manipulation of iSPNs, which bidirectionally altered lick responses in a manner consistent with their functional role in mediating inhibitory control over sound-driven action. Finally, Nrxn1α mutant mice exhibited corticostriatal synaptic deficits onto iSPNs in the TS, while demonstrating an impaired ability to withhold responses to non-target stimuli consistent with the role we uncovered for TS iSPNs in mediating response control.

### Both pathways promote behavioral responses to reward-associated auditory cues

Our photometry recordings revealed robust, concurrent activation of both striatal subtypes during stimulus presentation of the reward-predictive target stimulus in hit trials. Surprisingly, independent bilateral inhibition of either dSPNs or iSPNs was not sufficient to disrupt responding to the Go signal. Only simultaneous inhibition of both pathways using muscimol effectively impaired reward-related responses. Notably, this reduction in responding did not reflect a general motivational deficit or an inability to perform licking movements, as non-operant consummatory licking remained intact following muscimol administration. Instead, these findings suggest that broad inhibition of both striatal pathways in the TS selectively disrupts cue-response associations critical for reward-related behavior.

These results suggest that, alongside a reward-related dSPN component, iSPN ensembles in the TS may also contribute to generating responses to rewarded stimuli. This is consistent with work in other striatal areas demonstrating that concurrent activation of both striatal pathways is necessary for the initiation of actions^69–71^. Moreover, widely accepted models of striatal function propose that concurrently active iSPNs contribute to action control by suppressing alternative actions or by mediating the required timing and synchrony of striatal activity to initiate action^69,72–76^. Furthermore, recent single-unit recordings in the TS have identified a minority iSPN subpopulation that functionally aligns with dSPN-mediated auditory choices in a 2-AFC task^59^. Although single-unit recordings are needed to definitively characterize functional iSPN ensembles in this task, it is possible that a response-initiating iSPN subpopulation promotes action by laterally inhibiting distinct response-suppressing iSPN ensembles. This would be consistent with prior evidence that iSPNs serve as key mediators of lateral inhibition to both striatal pathways^21,24,26^. Future studies using cellular-resolution recording techniques will be critical for elucidating how the coordinated activity across striatal pathways in the tail mediates responses to reward-associated stimuli.

### TS iSPNs exhibit a task-related, enhanced baseline activity prior to stimulus presentation

We observed enhanced iSPN recruitment before sound presentation in both target and non-target trials. Although this tonic pattern of activity in iSPNs preceding sound onset was reduced in false alarm trials, further investigation didn’t support a role for this activity in directly mediating inhibitory control. Firstly, this activity was not continuously present during the no-lick period preceding trial onset, suggesting it does not mediate a general suppression of licking behavior. Furthermore, this tonic iSPN activity is not specific to trials requiring inhibitory control over sound driven behavior as it seems to emerge in the earliest stages of training before animals face operant licking contingencies. Finally, although we saw trends for the relationship of iSPN baseline and NoGo performance, the best predictor of inhibitory performance was the absolute peak of iSPN activity during sound presentation. Together these data suggest that while increased iSPN baseline activity may support inhibitory control, it does not appear to be sufficient for this process. Additionally, recent work in the TS has identified a similar pattern of iSPN activity prior to stimulus onset^77^. Tsuitsui-Kimura *et al*. observed enhanced baseline iSPN activity in the tail during trials in which mice were able to successfully overcome a threatening stimulus to gain reward. These data further suggest that persistent iSPN activity in TS does not necessarily serve to inhibit general movement.

Instead, the collective evidence suggests that rather than directly mediating inhibitory control, this tonic iSPN activity may reflect anticipatory signaling of incoming sensory cues relevant for guiding behavior. In line with this, both Tsuitsui-Kimura *et al*. and our work show this activity emerging throughout training when animals are learning to anticipate incoming sensory information. Furthermore, this interpretation may also explain why the absence of this activity correlates with erroneous performance in trained mice, as a lack of active sensory anticipation during such trials may lead them to revert to a more reactive mode of responding. If this is the case, it is possible that this persistent iSPN activity may be orchestrated by PFC projections that are sent to the TS^13,14,78–80^, supporting a top-down influence that interacts with the sensory-driven processing ongoing within the tail. This is consistent with our photometry findings, which show that neither baseline activity nor sound-evoked responses alone reliably predict NoGo performance. Rather, it is the aggregated contribution of both mechanisms that best accounts for behavioral performance in non-target trials. Future investigations into this striatal region should aim to understand the relative contributions of different cortical projections and how their integration shapes behavioral control of sensory-driven action.

### iSPNs in the TS contribute to inhibitory control over sensory-driven action

In addition to enhanced iSPN activity prior to sound onset, we observed greater recruitment of iSPNs compared to dSPNs during NoGo stimulus presentation in correct rejection trials. Moreover, our optogenetic inhibition of iSPNs during sound presentation supports a role for iSPNs in inhibitory control of sound-driven responses to inappropriate or unfavorable sensory stimuli. These findings are in line with previous reports in other striatal regions associating iSPN activity with the inhibition of unrewarded actions and subsequent facilitation of exploratory behavior^81,82^. Moreover, these cell-type specific manipulations extend previous work in non-human primates, implicating that GPe projecting neurons from the caudate tail are involved in the rejection of low-value visual stimuli^40,41^. Together, these data support proposed evolutionary homologies between the rodent TS and the primate caudate tail^32–34^, and provide mechanistic insight into indirect pathway involvement in inhibitory control, with future work aimed at evaluating the potential contributions of lateral inhibition within the TS to this process.

Importantly, we also demonstrated that manipulation of iSPNs in naïve mice does not bias free-licking behavior, supporting previous work proposing that this striatal region does not directly mediate general motor execution but rather contributes to perceptual decision-making^26,37,83^. Furthermore, recent work has proposed a particular involvement of the TS in facilitating sound-action associations, with dopamine signaling correlating to the reliability of a given action in a specific sensory context^84^. This is further consistent with literature implicating the TS in fear conditioning, during which a previously neutral stimulus comes to elicit a conditioned freezing response^32,77,85^. Our findings that iSPNs are sufficient to bias free-licking behavior in the absence of auditory stimuli, but only after animals have been trained to organize licking as a response to the Go stimulus, support a role for the TS in regulating learned stimulus-action associations. This interpretation is further strengthened by our muscimol experiments, in which inhibition of the TS selectively disrupted the association between the Go stimulus and the licking response.

Taken together, our optogenetic manipulations of iSPN activity reveals a clear role for these neurons in governing inhibitory control over learned sound-driven actions.

### Disruption of corticostriatal inputs onto iSPNs may underly impairments in inhibitory control in Nrxn1α-associated disorders

Another important finding of this study is the identification of a Nrxn1α-related synaptic deficit in the corticostriatal inputs onto iSPNs in the TS. This was consistent with recent work demonstrating impaired synaptic strength in corticostriatal projections of the PFC to the more anterior dorsomedial striatum in both Nrxn1α^+/−^ and Nrxn1α^-/-^ mice^45^. These data imply that impaired excitatory synaptic iSPN recruitment is a hallmark of the striatum of mice with Nrxn1α disruptions, a circuit vulnerability that could lead to broadly dysregulated cortical activity through the basal ganglia loop architecture. A limitation of the Nex-Cre mediated optogenetic strategy is our inability to determine the specific cortical inputs most impacted by the loss of Nrxn1α function. The extent and specificity of Nrxn1α associated synaptic deficits may differ across distinct cortical projection populations, an important consideration when parsing the contributions of PFC and auditory cortical dysregulation to altered behavior.

Interestingly, we found that Nrxn1α^-/-^ mice showed a selective deficit in inhibitory control over non-target stimuli, distinct from a general motor hyperactivity phenotype. Given the functional importance of TS iSPN activity identified in our study, it is likely that this synaptic deficit contributes to the observed learning impairments. Further supporting our observed behavioral deficit, similar NoGo learning impairments in Nrxn1α^-/-^ mice training in a visual Go/NoGo paradigm have recently been reported in a publicly available doctoral dissertation, lending convergent validity to our findings across independent laboratories and sensory modalities^86^. This observed behavioral phenotype is in line with the clinical characterization of Nrxn1α associated disorders, as impairments in sensory processing, sensorimotor gating, and inhibitory control are clinically relevant features in ASD, ADHD, Tourette’s Syndrome, and schizophrenia^86–92^. Furthermore, many of these disorders are highly comorbid with each other, suggesting commonalities in the underlying neural aberrations that give rise to these shared symptoms^89,93–95^. Given the established association of Nrxn1α copy number variants with these neurodevelopmental disorders, the circuit mechanisms identified in this study are likely to have clinical relevance and may represent a promising target for future therapeutic development.

## Materials and Methods

### Animals

All procedures and experiments were conducted in accordance with the National Institutes of Health Guidelines for the Use of Animals and approved by the University of Pennsylvania Institutional Animal Care and Use Committee (Protocol: 805643). All animals used in this study were adult mice (2.5-5 months). We used wild-type mice (C57BL/6J; Jackson Laboratory Strain: 000664), heterozygous mice bred from the Adora2a-Cre line (B6.FVB(Cg)-Tg(Adora2a-cre)KG139Gsat/Mmucd), and heterozygous mice bred from the Drd1a-Cre line (B6;129-Tg(Drd1-cre)120Mxu/Mmjax; MMRRC_037156-JAX). Homozygous Ai40D male breeders (B6.Cg-Gt(ROSA)26Sortm40.1(CAG-aop3/EGFP)Hze/J; IMSR_JAX:021188) were obtained from Jackson Laboratories to cross with either heterozygous Adora2a-Cre or Drd1a-Cre mice to generate double-transgenic mice used for experiments. From these breedings, we used Adora2a-Cre heterozygous; Ai40d heterozygous double-transgenic mice and Drd1a-Cre heterozygous; Ai40d heterozygous double-transgenic mice. Constitutive Nrxn1α KO mice were originally obtained from the Südhof lab and were maintained as previously described^45,96^. For slice experiments, Nrxn1α^+/−^ breeders were crossed to Nex^Cre/Cre^;R26R^AI32/AI32^;D1-Tomato mice (Nex-Cre mice: obtained with permission from Klaus-Armin Nave from the Zhou Lab, University of Pennsylvania; AI32:B6;129S-Gt(ROSA)26Sor-CAG-ChR2(H134R)-EYFP, JAX012569; D1-Tom: B6.Cg-Tg (Drd1a-tdTomato, JAX016204) to generate Nrxn1α^+/+^;Nex^Cre/+^;R26R^AI32/+^;D1^Tom^ or Nrxn1α^+/−^;Nex^Cre/+^;R26R^AI32/+^;D1^Tom^ experimental mice. Control animals for all experiments consisted of age– and sex-matched mixed littermates. All animals were housed with a 12:12 light-dark cycle with food provided ad libitum, and a restricted water schedule. Animals were typically group housed (2–5 per cage), with the exception of muscimol-treated cohorts and individuals displaying aggressive behavior towards cage mates.

### Behavioral Apparatus and Training

To assess value-based responses to auditory stimuli we employed a head-fixed auditory Go/NoGo behavior where licking was used to register a response. Behavioral apparatuses were custom-built within double-walled sound-attenuating chambers containing head-fixation clamps (ThorLabs), a 3-D printed platform (derived from a treadmill design by Janelia J005558), an optical lickometer (Sanworks 1020), trial start LED (either white or green as it served as a mask for optogenetic experiments), water dispensing system and sound system. Behavioral programs were operated using custom built circuits with Arduino Uno and Teensy 3.2 microcontrollers and custom software. Behavioral data were acquired via serial communication and saved as CSV files using the CoolTerm terminal application.

Our water dispensing system consisted of a solenoid valve (The Lee Company LHDA1231115H) with tubing (Tygon S3 E-3603) and an elevated water reservoir (30mL syringe; Fisher Scientific 22124969). As water reward amount depended on the duration the solenoid valve was open and the height of the reservoir, the system was regularly calibrated to reliably dispense 4uL reward volumes. Furthermore, water was replenished to the same height at the start of each session to maintain consistent fluid pressure.

Our sound system included an Adafruit Audio FX Sound Board (16MB) connected to output speakers (DigiKey 668-1447-ND) for playback of bandlimited sound stimuli. These stimuli (5–7.2 kHz and 13.3–19.2 kHz) were generated using custom MATLAB code by applying calibrated bandpass filters to white noise, ensuring flattened spectral profiles across the target frequency ranges. To prevent spectral artifacts, all stimuli were onset– and offset-ramped using 5 ms raised cosine (cosine-squared) envelopes.

Prior to training, mice were water restricted to >85% of their initial body weight over the course of 2-3 days during which animals became familiar with experimental handling. Next, animals underwent 1-2 sessions of magazine training where mice acclimated to head-fixation in the box by freely receiving reward. Animals then underwent 2 stages of behavioral shaping before reaching final Go/NoGo training. Cohorts were counterbalanced in which stimulus was assigned as the Go versus NoGo cue for training. For all stages, each training session was ∼30 minutes long and every trial began with a no-lick, dark period of 3 consecutive seconds before the box light turned on.

During initial Association Training, the Go sound was played for 1 second immediately before a randomly delivered reward (occurring 1-10s after trial start). Animals were considered experts in this stage after training for 10 days, or after receiving >2RW/min and demonstrating a significant increase in anticipatory licking during the cue compared to 1s prior (for two consecutive days).

During the Go Training stage, animals learned to lick within 1s of the end of a 150ms Go sound (total response window being 1.15s). During this stage they also learned to withhold premature licking prior to the sound presentation. From this phase onward, a premature lick resulted in a 5s lockout penalty. For initial Go training, the delay from trial start to Go sound presentation ranged from 0.5-1s and misses (defined as a failure to lick within 1s after offset of the Go sound) were not punished. Once animals achieved a d’≥1.5 (calculated by comparing the hit rate and premature rate) for a session, animals progressed to the final Go training phase where the sound delay increased to up to 2s and misses were punished with a 5s lockout penalty. Animals were considered experts at this stage once they reached a d’≥2 for 3 consecutive sessions.

Finally, during Go/NoGo training NoGo trials appeared randomly at a 40% probability during which animals received a 5s lockout penalty for false alarm responses but avoided this penalty when correctly withholding their licking to the NoGo stimulus.

Animals were considered to be “experts” after reaching a d’≥2 for 3 consecutive sessions. During training sessions, incorrect trials were repeated, however during all behavioral manipulation sessions trials were presented in a fully random order.

### Behavioral Analysis

For analysis of behavioral data we employed custom MATLAB scripts. To quantify Go/NoGo performance we used standard signal detection theory metrics. Discrimination index or d’ was quantified as d′ = Z(Hit Rate) – Z(False Alarm Rate) for the Go/NoGo stage and d′ = Z(Hit Rate) – Z(Premature Rate) for Go training stage. Response bias or the criterion metric was quantified as c =−0.5[Z(Hit Rate)+Z(False Alarm Rate)]. As animals often had hit rates at 1 for a given session, all rates that were at 0 or 1 extremes were adjusted relative to the total number of Go or NoGo trials. As our paradigm does not require the animal to initiate trials, engagement was quantified during analysis by tracking the local miss rate and lick rate following reward. If the local miss rate rose past the engagement cutoff (or if animals stopped ingesting reward) the remainder of the trials for that session were not analyzed. Engagement cutoffs were only used to calculate performance during training as well-trained animals typically maintained their engagement throughout the entire 30-minute behavioral sessions.

### Surgical procedures

All surgical procedures were performed on a stereotaxic frame (Kopf Instruments, Model 1900). Briefly, mice were anesthetized using vaporized isoflurane (1-2% + oxygen at 1.5 L/min). Body temperature was continuously monitored and maintained at 30°C during surgery (Harvard apparatus, #50722F; 55-7030). After administration of ophthalmic ointment (Puralube Vet Ointment) and analgesics (Meloxicam-SR; bupivacaine), the scalp was prepared for surgery by removing overlying fur with depilatory cream, followed by alternating washes with 70% ethanol and betadine. A small anterior– posterior incision was made using a scalpel to expose the skull, which was then crosshatched with scalpel marks for texturization and cleaned with hydrogen peroxide. Bilateral craniotomies for injections and fiber implantations were made by drilling small (0.5mm) holes above the target coordinates for the TS (AP: +2.15-2.2mm from the interaural line (–1.7mm from bregma), ML: +/− 3.5mm, DV: –3.0mm from cortical surface for injections, –2.6 to –2.8mm for fiber placements).

Injections were performed using the Nanoject II/III system and a pulled glass needle backfilled with mineral oil. A total volume of 350–368 nL of virus was injected at a rate of ∼1 nL/sec per injection site. Following each injection, the glass needle remained in place for about 10-15mins before being slowly withdrawn out to minimize backflow. For optogenetic excitation experiments we used AAV8-hSyn-FLEX-ChrimsonR-tdTomato (UNC Vector Core) at a titre of 3.7*10^12^ V.G./mL). For photometry experiments, we used pGP-AAV9-syn-FLEX-jGCaMP8m-WPRE (Addgene) with an original titre of 2.4*10^13^ V.G./mL diluted to 4.8*10^12^ V.G./mL.

For all optogenetic experiments, we used fiber optic cannulae (RWD; R-FOC-BL200C-50NA) with white/black Φ1.25mm ceramic ferrules with 0.5ΝΑ, 200μm fiber optic cores at a length of 4mm. For all photometry experiments we used fiber optic cannulae (RWD; R-FOC-BL200C-39NA) with black Φ1.25mm ceramic ferrules with 0.39ΝΑ, 200μm fiber optic cores at a length of 4mm. Fibers were slowly lowered into the open craniotomies and secured in place with dental cement (Parkell C&B Metabond Quick Adhesive Cement System). Custom-made stainless steel headplates (Perelman Research Instrumentation Shop; eMachineShop) were also secured to the skull using Metabond with the remainder of the headcap being constructed with clear or black tinted Ortho-Jet dental acrylic (Lang Dental 1320CLR; 1306CLR; 3302BLK).

### Fiber photometry recordings

Batched GCaMP fluorescence was recorded using the Neurophotometrics fiber photometry system (FP3002; MBF Bioscience) while animals performed the behavioral task. Briefly, we used time-dependent modulation in which 415 nm and 470 nm LEDs were alternated at a combined frequency of 80 Hz (40 Hz per wavelength) to capture isosbestic and calcium-dependent GCaMP fluorescence, respectively. Emission was collected via a 200 μm core, 0.39 NA fiber optic patch cord and detected by an internal CMOS camera with sub-millisecond synchronization. Excitation power at the patch cord tip was calibrated to 20–40 μW per wavelength using an external power meter. Recordings were acquired and initially processed using Bonsai, with behavioral events time-locked to the fluorescence signal via 5V TTL input signals delivered from an Arduino Uno.

### Photometry analysis

Fluorescence signals were preprocessed using a custom MATLAB pipeline adapted from previously published code^97^. Raw data were separated by channel (415 nm isosbestic and 470 nm GCaMP), and the first 400 frames were discarded to remove initial instability. Signals were then low-pass filtered at 5 Hz (4th-order Butterworth) to reduce high-frequency noise. Slow drift due to photobleaching was corrected by fitting a cubic polynomial to each channel and computing ΔF/F as the residual divided by the fitted curve. To isolate calcium-dependent fluorescence changes, we performed a robust linear regression of the debleached isosbestic ΔF/F signal onto the debleached GCaMP ΔF/F signal. The fitted isosbestic component was then subtracted from the GCaMP trace to remove shared artifacts, and the resulting artifact-corrected ΔF/F trace was z-scored across the entire session.

To obtain group fluorescence traces, the z-scored ΔF/F (zΔF/F) values surrounding the event (typically sound onset) were averaged across trials to obtain an average fluorescence trace for each recording site. These average traces were then averaged across recording sites to obtain the group fluorescence trace. For quantification of baseline fluorescence preceding trial epochs, zΔF/F fluorescence values from the 500ms immediately before the event were averaged for each trial. The average baseline fluorescence across trials was then obtained for each recording site. To quantify the absolute peak fluorescence values, the maximum zΔF/F values in the 375ms window following the trial event was obtained for each trial. These values were then averaged across trials to obtain the average absolute peak fluorescence for each recording site. To obtain the sound-evoked peak fluorescence (i.e. peak prominence or baseline subtracted peak), the baseline fluorescence value for each trial was subtracted from the corresponding absolute peak value. These baseline-subtracted peaks were then averaged across trials to obtain the sound-evoked peak fluorescence values for each recording site.

To quantify relationships between neural activity and behavioral performance, we used linear mixed-effects models (LMMs) implemented in MATLAB using the fitlme function. Models were constructed with the average sound-evoked photometry responses for recording site as the dependent variable and behavioral measures such as d′ or response bias as fixed effects. To account for repeated measures across recording sites, we included a random intercept for each site, modeling the within-subject correlation structure across sessions. Model assumptions were checked by inspecting residuals, and statistical significance of fixed effects was evaluated using t-statistics and corresponding p-values.

### Optogenetic manipulations

To mask effects of the stimulation light for all optogenetic experiments, we used an LED placed near the mouse’s visual field which also served as the trial start light. For bilateral optogenetic excitation with ChrimsonR, we used ∼625nm LED light sources (Prizmatix Fiber Coupled Dual Optogenetic-LED; Orange-Red) connected to a calibrated to an output of about 5mW as measured at the tip of the splitter patch cord (Doric Splitter Branching Patch Cord; SBP(2)_200/230/900-0.57_1m_FCM-2xMF1.25).

For optogenetic manipulations during the Go/NoGo task, we used an Arduino with custom software to deliver 250ms of 20Hz pulsed light time-locked to sound presentation, beginning 50 ms before sound onset and ending 50 ms after sound offset on a randomized 25% of trials. All excitatory stimulation experiments used 20Hz pulsed light to minimize channel desensitization.

For bilateral optogenetic inhibition with ArchT, we used 532nm laser light sources (Shanghai Dream Lasers Technology; DPSS, SDL-532-100T) connected to a calibrated to an output of about 10mW as measured at the tip of the splitter patch cord. For optogenetic manipulations during the Go/NoGo task, we used a similar stimulation protocol as described for excitation except we used continuous illumination. For each experiment, all animals underwent at least 3 sessions of opto during Go/NoGo and sessions with a d’ of less than 2 were excluded from analysis. All of the trials from these sessions were then aggregated to calculate the final behavioral metrics based on trial type.

To examine the effects of optogenetic manipulation of SPNs outside of our direct behavioral context, we ran mice in a non-operant session within the same behavioral boxes, during which licking had no impact on water availability but was being continuously recorded. In these non-operant control sessions, no sounds were presented throughout the duration of the session, which consisted of at least 100 stimulation trials. The box was illuminated the entire duration of the session. Throughout the session, blocks of 10 stimulation trials (in which the stimulation LED was turned on for 1s) or 10 control trials (in which the mask light was activated) were presented. These blocks were interleaved with water blocks, during which the animal received 5 randomly dispensed water rewards to encourage free licking. Water delivery, stimulation, and mask light illumination occurred at a randomized interval of 1–10 seconds following trial onset. Free licking rates were calculated for each stimulation trial in the 1s prior to stimulation, 1s during stimulation, and 1s after stimulation. These rates were then averaged to obtain the average lick rate for each animal during each respective bin.

### Muscimol Inactivation

Pharmacological inhibition using muscimol was performed as previously described^37,38^. Incomplete bilateral craniotomies (0.5mm diameter) were positioned dorsally over the TS and labelled with permanent ink during initial head plating surgeries. The head plate was stereotactically positioned during this surgery and bregma remained exposed to allow for accurate injection targeting later during head fixation in our customized injection stereotax. The skull was then covered with Kwik-Sil (World Precision Instruments), and animals were singled housed to prevent exposure of the skull. Immediately prior to injections, animals were lightly anesthetized using isoflurane prior to head fixation in the stereotax. While under isoflurane, body temperature was maintained using a 30°C warming pad (Kent Scientific, Far Infrared Warming Pad with Controller (15.2 cm W x 20.3 cm L)). The Kwik-Sil was then removed, and the craniotomies were fully opened. Injections were performed using a Nanoject II (Drummond) fitted with a pulled glass pipette, delivering 36.8nL to each hemisphere at a rate of ∼0.6 nL/s. Either PBS vehicle or 0.5 mM muscimol (Sigma-Aldrich, M1523) was co-injected with 0.1mg/mL Alexa Fluor 647-conjugated cholera toxin subunit B (Fisher Scientific, C34778) to enable histological verification of injection sites. Final muscimol dose per hemisphere was roughly 2ng. Each injection was allowed to sit for 60s to minimize any backflow before slowly removing the injection pipette. Following the final injection, the injection sites were protected with saline and Kwik-Sil and animals were returned to their home cage which was placed on a heating pad (K&H Pet Products Small Animal Outdoor Heated Pad) to allow for 30 minutes of recovery time prior to behavioral testing. Following recovery, mice were run for in a non-operant version of the behavior where licking had no impact on reward availability. In this session, animals were able to lick freely and water was dispensed following a randomized 1s presentation of the Go stimulus. After about 5 mins (or 40 rewards) the program was switched to the actual Go/NoGo paradigm where mice were run for 30 mins. On muscimol injection days, mice ran in additional washout sessions after the effects of the acute injection had largely ceased.

### Electrophysiology

Acute slice electrophysiology experiments were performed as previously described^45,98^. In brief, mice were anesthetized and perfused using aCSF containing (in mM): 124 NaCl, 2.5 KCl, 1.2 NaH2PO4, 24 NaHCO3, 5 HEPES, 12.5 Glucose, 1.3 MgSO4, 2.5 CaCl2. Following dissection, the brain was submerged in ice-cold aCSF and 250μm coronal sections were made using a vibratome (Leica VT1200). Slices were then transferred to a 32°C NMDG recovery solution comprised of (in mM): 92 NMDG, 2.5 KCl, 1.2 NaH2PO4, 30 NaHCO3, 20 HEPES, 25 Glucose, 5 Sodium ascorbate, 2 Thiourea, 3 Sodium pyruvate, 10 MgSO4, 0.5 CaCl2. Following 12–15 min recovery, slices were then transferred to room temperature aCSF chamber (20–22°C) and left for at least 1 h before recording. During recordings, slices were fully submerged in an oxygenated (95% O2, 5% CO2) 29–30°C aCSF containing Picrotoxin (100 uM, Hello Bio), released at a flow rate of 1.4–1.6 mL/min.

For voltage-clamp recordings, recording pipettes were pulled from borosilicate glass (World Precision Instruments, TW150-3) with a tip resistance of 3-5 MΩ when filled with internal solution containing (in mM) 115 CsMeSO3, 20 CsCl, 10 HEPES, 0.6 EGTA, 2.5 MgCl, 10 Na-Phosphocreatine, 4 Na-ATP, 4 Na-GTP, 0.1 Spermine, 1 QX-314 (pH adjusted to 7.3-7.4 with CsOH). To optogeneticially evoke cortical inputs, we used a full-field 470nm illumination from a collimated LED illuminator (CoolLED, PE-300) through a 40x objective (Olympus, 0.8NA water immersion) with a pulse width of 1ms. Optical-evoked voltage-clamp recordings (v_hold_ = –80mV) were performed in the presence of picrotoxin (100 μM), a GABAA antagonist. LED intensities ranged from 0.57-1.65 mW/mm2 during the optical input output measurements.

Paired-pulse ratios (PPR) were measured by optically evoking AMPA receptor– mediated excitatory postsynaptic currents (EPSCs). Each paired-pulse trial consisted of two light pulses of equal intensity and duration, applied at inter-stimulus intervals (ISIs) of 50, 100, 200, or 500 ms. All recordings were performed in the continuous presence of picrotoxin (PTX, 100 μM) to isolate glutamatergic currents. For each ISI, 5 trials were collected from each cell, and successive sets of trials were separated by a 10 sec interval. The amplitude of the second EPSC (EPSC2) was normalized to the amplitude of the first EPSC (EPSC1) to calculate the PPR (EPSC2/EPSC1).

## Supplemental Figures

**Figure S1.**
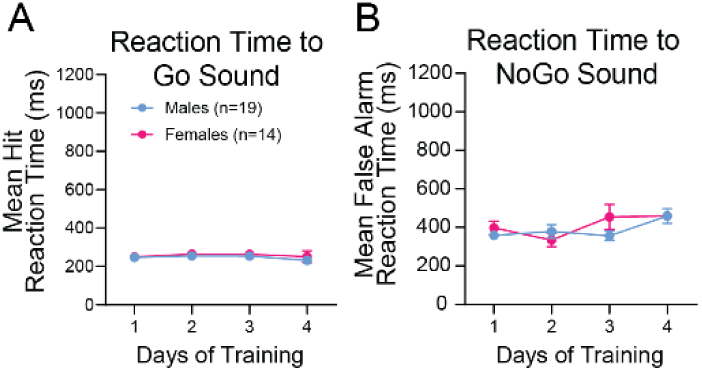
Supplementary behavior figure. (A) Reaction times to Go sound across Go/NoGo training for male (n = 19) and female (n = 14) mice. Symbols signify group means ± SEM. Linear mixed effects model with training day and sex as fixed effects, and subject as a random effect. There were no significant main effects of either training day (F(1.027, 23.96) = 1.74, p = 0.1995; Geisser-Greenhouse correction applied) or sex (F(1, 31) = 0.33, p = 0.57) for hit reaction times. (B) Reaction times to NoGo sound across Go/NoGo training for male (n = 19) and female (n = 14) mice. Symbols signify group means ± SEM. Linear mixed effects model with training day and sex as fixed effects, and subject as a random effect. There were no significant main effects of either training day (F (2.234, 52.13) = 1.031, p = 0.37; Geisser-Greenhouse correction applied) or sex (F (1, 31) = 0.42, p = 0.52) for false alarm reaction times.

**Figure S2.**
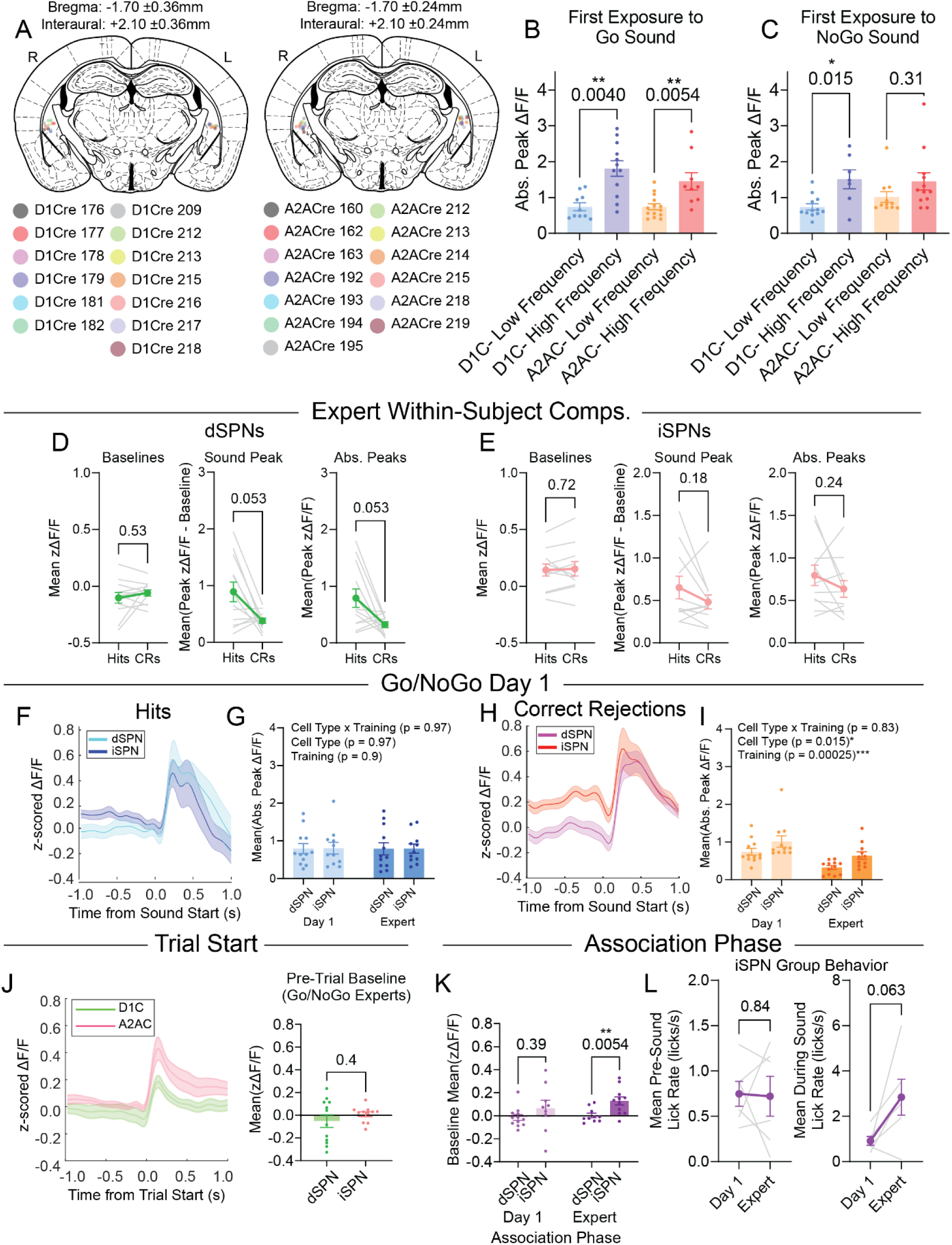
Photometry supplement 1. (A) Summary fiber placements for (left) D1Cre and (right) A2ACre photometry mice. (B) Mean absolute peak fluorescence values during the first day of association training for each cell-type and the assigned Go frequency. Dots signify individual recording sites [D1C: n_low_ = 6 mice/10 sites, n_high_ = 6 mice/12 sites; A2AC: n_low_ = 7 mice/14 sites, n_high_ = 5 mice/9 sites] with bars denoting group mean ± SEM. Comparisons across D1C mice were performed using a two-way mixed-effects model (REML) with Go Frequency and Hemisphere as fixed effects and Subject as a random effect; Go Frequency, p = 0.0040**; Hemisphere, p = 0.2752; interaction, p = 0.6596. Comparisons across A2AC mice, two-way mixed-effects model (REML) with Go Frequency and Hemisphere as fixed effects and Subject as a random effect; Go Frequency, p = 0.0054**; Hemisphere, p = 0.8869; interaction, p = 0.3099. (C) Mean absolute peak fluorescence values during correct rejection trials on the first day of Go/NoGo training for each cell-type and the assigned NoGo frequency. Dots signify individual recording sites [D1C: n_low_ = 6 mice/12 sites, n_high_ = 5 mice/7 sites; A2AC: n_low_ = 6 mice/11 sites, n_high_ = 6 mice/12 sites] with bars denoting group mean ± SEM. Comparisons across D1C mice were performed using a two-way mixed-effects model (REML) with NoGo Frequency and Hemisphere as fixed effects and Subject as a random effect; NoGo Frequency, p = 0.0148*; Hemisphere, p = 0.0129*; interaction, p = 0.1996. Comparisons across A2AC mice, two-way mixed-effects model (REML) with NoGo Frequency and Hemisphere as fixed effects and Subject as a random effect; NoGo Frequency, p = 0.3106; Hemisphere, p = 0.0832; interaction, p = 0.6916. (D) dSPN baseline and peak fluorescence comparisons between hit and correct rejection trials within a single expert session (n = 6 mice/12 sites). All analyses performed using a two-way mixed-effects model (REML) with trial type and hemisphere as fixed effects and subject as a random effect. Baselines: trial type, p = 0.5251; Hemisphere, p = 0.7818; interaction, p = 0.5367. Baseline-subtracted peak: trial type, p = 0.0527; Hemisphere, p = 0.7701; interaction, p = 0.2763. Absolute peak: trial type, p = 0.0534; Hemisphere, p = 0.6887; interaction, p = 0.2958. (E) iSPN baseline and peak fluorescence comparisons between hit and correct rejection trials within a single expert session (n = 6 mice/11 sites). All analyses performed using a two-way mixed-effects model (REML) with trial type and hemisphere as fixed effects and subject as a random effect. Baselines: trial type, p = 0.7153; Hemisphere, p = 0.5820; interaction, p = 0.7341. Baseline-subtracted peak: trial type, p = 0.1842; Hemisphere, p = 0.8547; interaction, p = 0.4585. Absolute peak: trial type, p = 0.2379; Hemisphere, p = 0.8809; interaction, p = 0.5351. (F) Mean fluorescence traces for hit trials during the first day of Go/NoGo training for dSPNs (n = 6 mice; 12 sites) and iSPNs (n = 6 mice, 11 sites). Shading denotes SEM. (G) Comparison of average hit absolute peak fluorescence for d/iSPNs during the first Go/NoGo session and expert session. Two-way mixed-effects model (REML) with pathway and training as fixed effects and recording site as a random effect. Pathway, p = 0.97; Training, p = 0.90; Interaction, p = 0.97. (H) Mean fluorescence traces for correct rejection trials during the first day of Go/NoGo training for dSPNs (n = 6 mice; 12 sites) and iSPNs (n = 6 mice, 11 sites). Shading denotes SEM. (I) Comparison of average correct rejection absolute peak fluorescence for d/iSPNs during the first Go/NoGo session and expert session. Two-way mixed-effects model (REML) with pathway and training as fixed effects and recording site as a random effect. Pathway, p = 0.015*; Training, p = 0.0002***; Interaction, p = 0.83. (J) Mean fluorescence traces aligned to trial start during an expert Go/NoGo session for dSPNs (n = 6 mice/12 sites) and iSPNs (n = 6 mice/11 sites). Shading denotes SEM. Pre-trial baseline did not significantly differ between dSPNs and iSPNs (mixed-effects model (REML); Pathway, p = 0.4045; Hemisphere, p = 0.2928; Interaction, p = 0.2272). (K) Comparison of average pre-sound baseline fluorescence for dSPNs and iSPNs during the first and expert association training sessions. All analyses performed using mixed-effects model (REML) with pathway and hemisphere as fixed effects and subject as a random effect. Day 1: Pathway, p = 0.3887; Hemisphere, p = 0.6462; Interaction, p = 0.1168 [N_D1C_ = 6 mice/12 sites; N_A2AC_ = 5 mice/9 sites]. Expert: Pathway, p = 0.0054**; Hemisphere, p = 0.5864; Interaction, p = 0.1226 [N_D1C_ = 5 mice/10 sites; N_A2AC_ = 6 mice/11 sites]. (L) Average lick rates for the mice in the iSPN cohort (n = 6) before sound presentation (left) and during sound presentation (right). Comparisons performed using wilcoxon matched-pairs signed rank test; Pre-sound (W = –3, p = 0.8438), During sound (W = 19, p = 0.0625).

**Figure S3.**
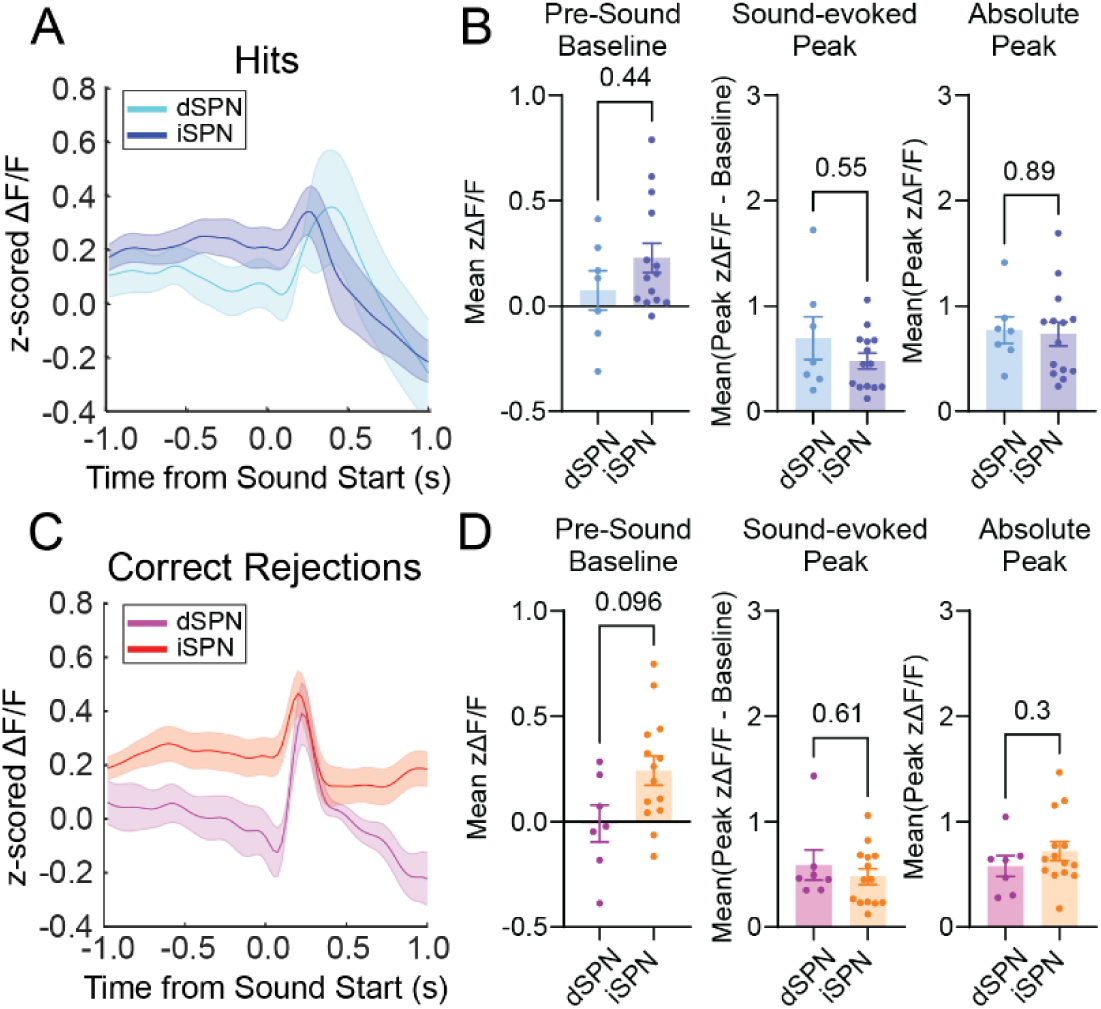
Low-frequency Go sound photometry cohort. (A) Average fluorescence traces for dSPNs (light blue; n = 5 mice/7 sites) and iSPNs (dark blue; n = 7 mice/14 sites) during hit trials in a Go/NoGo expert session. Shading denotes SEM. (B) Quantification for the baseline, sound-evoked peak, and absolute peak fluorescence values for d/iSPN expert hit trials. Mixed-effects model (REML) with Pathway and Hemisphere as fixed effects, and Subject as a random effect. Baseline: Pathway, p = 0.4392; Hemisphere, p = 0.8143; Interaction, p = 0.0528. Baseline-subtracted peak: Pathway, p = 0.5494; Hemisphere, p = 0.0268*; Interaction, p = 0.0388*. Absolute peak: Pathway, p = 0.8851; Hemisphere, p = 0.0804; Interaction, p = 0.7640. (C) Average fluorescence traces for dSPNs (purple; n = 5 mice/7 sites) and iSPNs (orange; n = 7 mice/14 sites) during correct rejection trials in a Go/NoGo expert session. Shading denotes SEM. (D) Quantification for the baseline, sound-evoked peak, and absolute peak fluorescence values for d/iSPN expert correct rejection trials. Mixed-effects model (REML) with Pathway and Hemisphere as fixed effects, and Subject as a random effect. Baseline: Pathway, p = 0.0958; Hemisphere, p = 0.9167; Interaction, p = 0.1320. Baseline-subtracted peak: Pathway, p = 0.6047; Hemisphere, p = 0.2153; Interaction, p = 0.3181. Absolute peak: Pathway, p = 0.3009; Hemisphere, p = 0.2649; Interaction, p = 0.8104.

**Figure S4.**
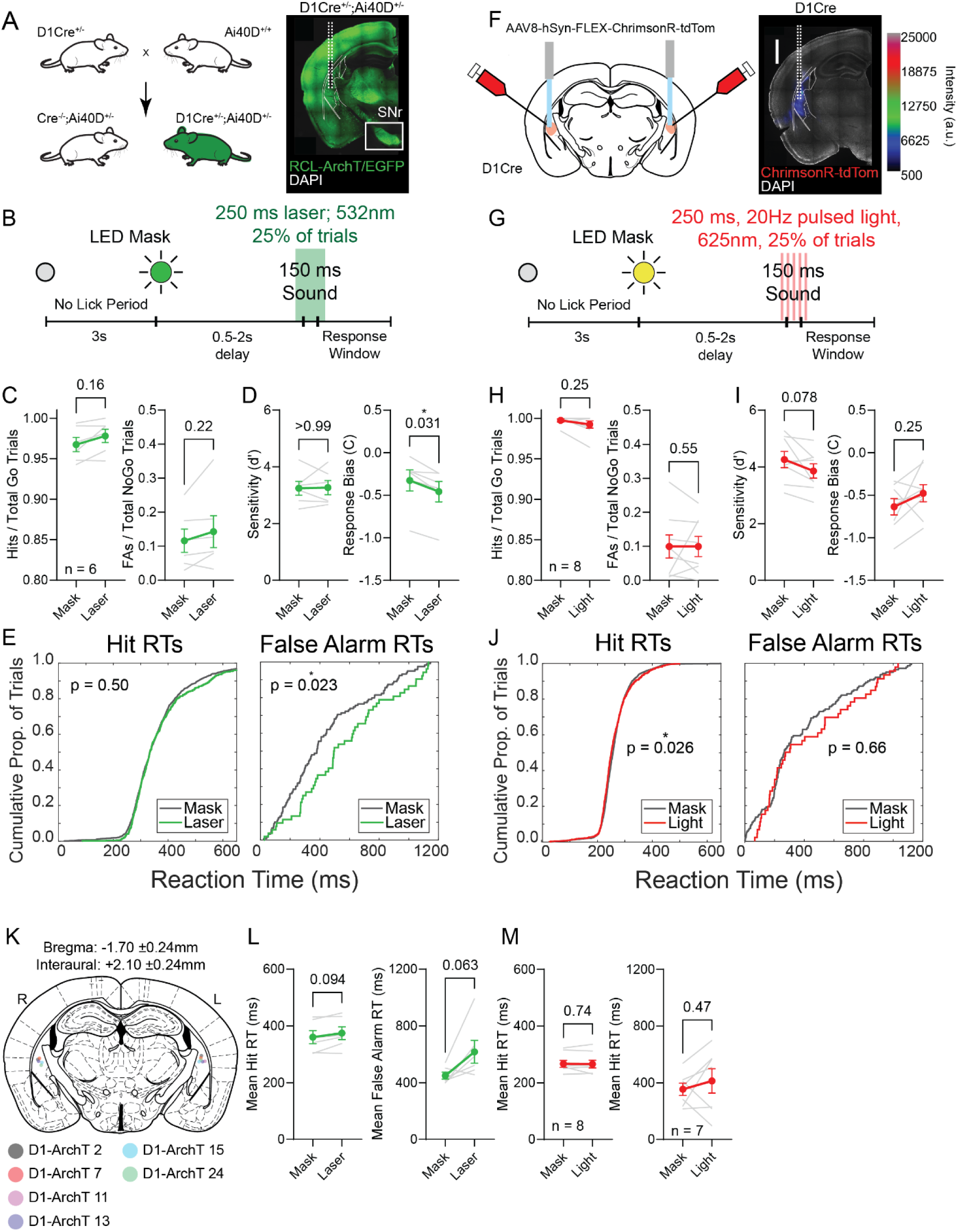
Manipulations of dSPNs alone have minimal impacts on behavior. (A) Experimental breeding strategy to generate transgenic mice expressing ArchT in dSPNs with example histology image. Inset shows dSPN terminal expression in the substantia nigra pars reticulata. (B) Schematic showing time course for laser-mediated inhibition of dSPNs during sound presentation. (C) Hit and false alarm rates for mask control versus laser mediated inhibition trials (n = 6). Two-tailed Wilcoxon matched-pairs signed-rank test (exact): Hits (W = 15, p = 0.1562), FAs (W = 13, p = 0.2188). (D) Discrimination sensitivity and response bias values for mask control versus laser mediated inhibition trials (n = 6). Two-tailed Wilcoxon matched-pairs signed-rank test (exact): d’ (W = 1, p > 0.9999), C (W = –21, p = 0.0312*). (E) Cumulative distributions for hit and false alarm reaction times (RT) during mask and laser inhibition conditions. Curves pool trials from multiple sessions across 6 mice [Mask: n_hit_ = 1627, n_FA_ = 119; Laser: n_hit_ = 539, n_FA_ = 52]. Differences between distributions were tested with two-sample Kolmogorov-Smirnov tests (two-sided): Hits, D = 0.0407, p = 0.5044; False Alarms, D = 0.2432, p = 0.0227*. Effect on false alarm RT was only a trend on subject-level comparison (see L). (F) Experimental schematic to expressing excitatory red-shifted opsin ChrimsonR in dSPNs with example histology image. Scale bar is 1mm. (G) Schematic showing time course for light-mediated excitation of dSPNs during sound presentation. (H) Hit and false alarm rates for mask control versus light mediated excitation trials (n = 8). Two-tailed Wilcoxon matched-pairs signed-rank test (exact): Hits (W = –8, p = 0.25), FAs (W = –10, p = 0.5469). (I) Discrimination sensitivity and response bias values for mask control versus light mediated excitation trials (n = 8). Two-tailed Wilcoxon matched-pairs signed-rank test (exact): d’ (W = –26, p = 0.0781), C (W = 18, p = 0.25). (J) Cumulative distributions for hit and false alarm reaction times (RT) during mask and light conditions in excitation experiment. Curves pool trials from multiple sessions across 6 mice [Mask: n_hit_ = 2267, n_FA_ = 140; Laser: n_hit_ =768, n_FA_ = 46]. Differences between distributions were tested with two-sample Kolmogorov-Smirnov tests (two-sided): Hits, D = 0.0612, p = 0.0261*; False Alarms, D = 0.1214, p = 0.6586. Effect on hit RT was not recapitulated with subject-level comparison (see M). (K) Summary fiber placements for D1-ArchT+ mice. (L) Average reaction time by animal for hits (left) and false alarm (right) trials across mask and laser mediated inhibition trials (n = 6). Two-tailed Wilcoxon matched-pairs signed-rank test (exact): Hits (W = 17, p = 0.0938), FAs (W = 19, p = 0.0625). (M)Average reaction time by animal for hits (left) and false alarm (right) trials across mask and light mediated excitation trials. One animal was omitted from the false alarm comparison because they did not have any false alarms for the light on condition. Two-tailed Wilcoxon matched-pairs signed-rank test (exact): Hits (n = 8, W = 6, p = 0.7422), FAs (n = 7, W = 10, p = 0.4688).

**Figure S5.**
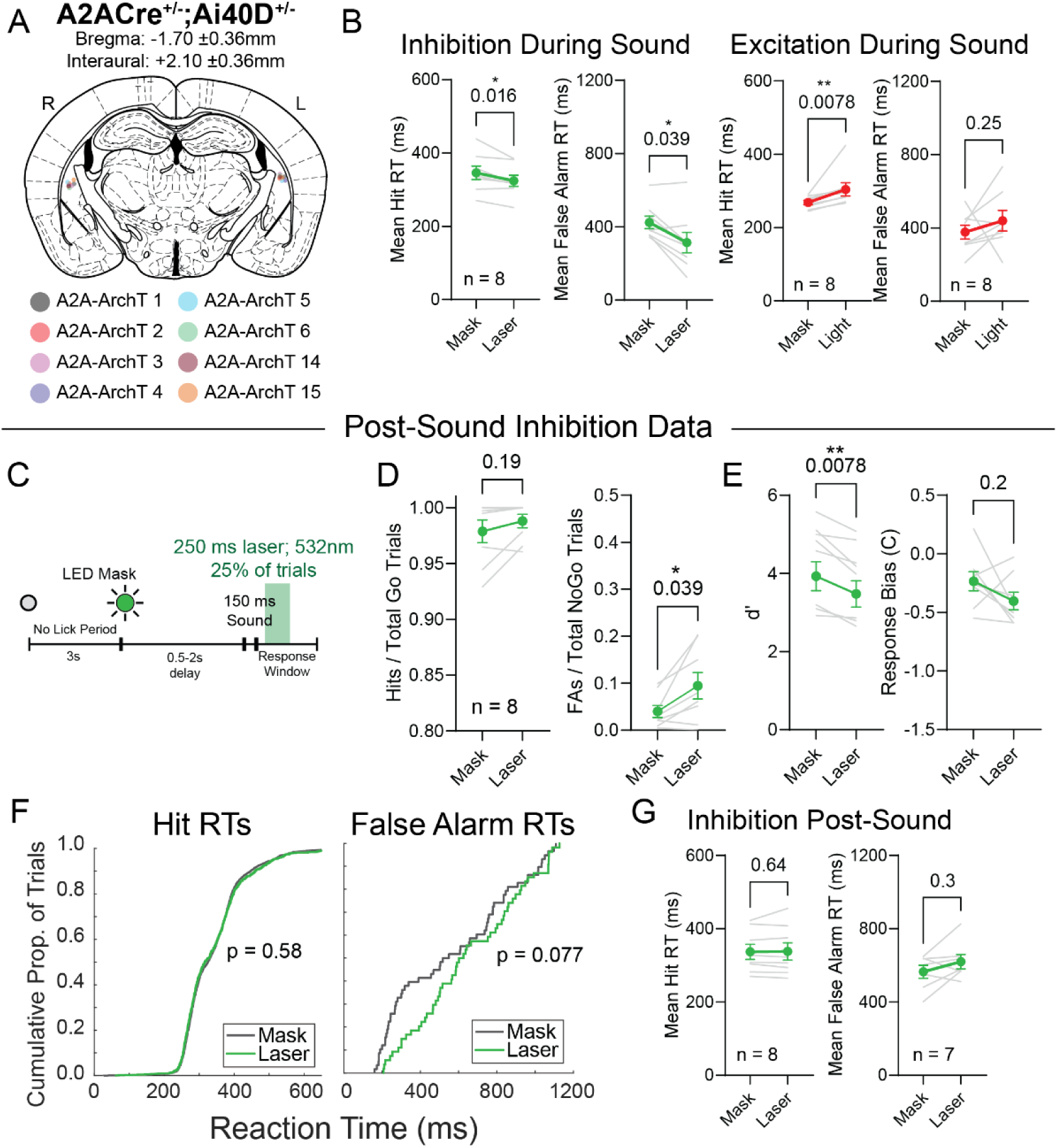
Post-sound inhibition of iSPNs. (A) Summary fiber placements for A2A-ArchT+ mice. (B) (left pair) Average reaction time by animal for hits and false alarm trials across mask and laser mediated sound inhibition trials (see Fig. 4E; n = 8). Two-tailed Wilcoxon matched-pairs signed-rank test (exact): Hits (W = –34, p = 0.0156*), FAs (W = –30, p = 0.0391*). (right pair) Average reaction time by animal for hits and false alarm trials across mask and light-mediated excitation trials (see Fig. 4J; n = 8). Two-tailed Wilcoxon matched-pairs signed-rank test (exact): Hits (W = 36, p = 0.0078**), FAs (W = 18, p = 0.25). (C) Schematic showing time course for laser-mediated inhibition of iSPNs after sound presentation. (D) Hit and false alarm rates for mask control versus laser mediated post-sound inhibition trials (n = 8). Two-tailed Wilcoxon matched-pairs signed-rank test (exact): Hits (W = –11, p = 0.1875), FAs (W = 30, p = 0.0391*). (E) Discrimination sensitivity and response bias values for mask control versus laser mediated post-sound inhibition trials (n = 8). Two-tailed Wilcoxon matched-pairs signed-rank test (exact): d’ (W = –36, p = 0.0078**), C (W = –20, p = 0.1953). (F) Cumulative distributions for hit and false alarm reaction times (RT) during mask and laser inhibition conditions. Curves pool trials from multiple sessions across 8 mice [Mask: n_hit_ = 2615, n_FA_ = 58; Laser: n_hit_ = 821, n_FA_ = 54]. Differences between distributions were tested with two-sample Kolmogorov-Smirnov tests (two-sided): Hits, D = 0.0309, p = 0.5844; False Alarms, D = 0.2350, p = 0.0771. (G) Average reaction time by animal for hits (left) and false alarm (right) trials across mask and laser mediated post-sound inhibition trials. One animal was excluded from the right panel because it did not have any FA trials during the laser condition. Two-tailed Wilcoxon matched-pairs signed-rank test (exact): Hits (n = 8, W = –8, p = 0.6406), FAs (n = 7, W = 14, p = 0.2969).

**Figure S6.**
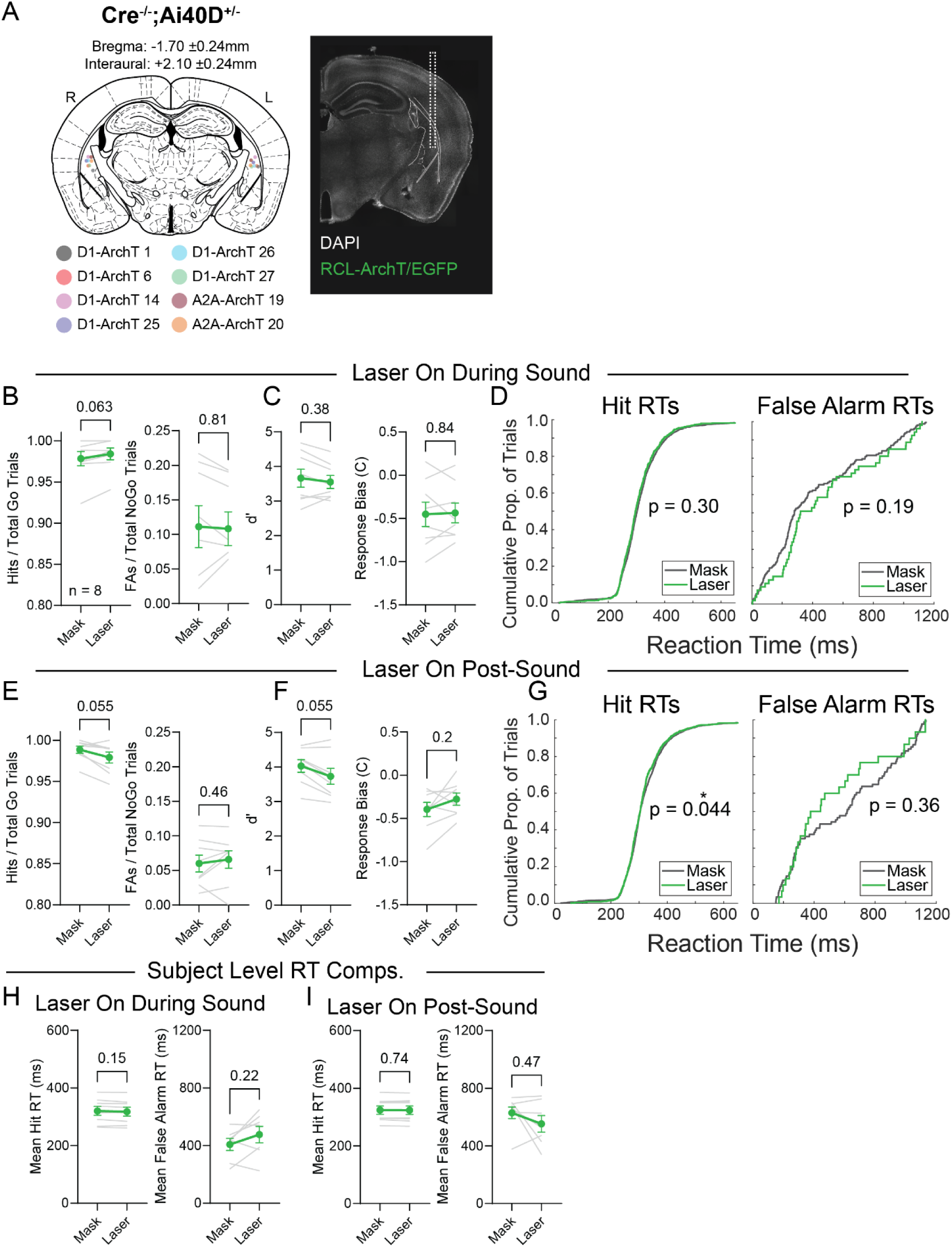
Cre-Negative mice do not display light-induced effects. (A) Summary fiber placements for Cre-Negative mice along with example histological image. (B) Hit and false alarm rates for mask control versus laser on during sound trials (n = 8). Two-tailed Wilcoxon matched-pairs signed-rank test (exact): Hits (W = 19, p = 0.0625), FAs (W = –4, p = 0.8125). (C) Discrimination sensitivity and response bias values for mask control versus laser on during sound trials (n = 8). Two-tailed Wilcoxon matched-pairs signed-rank test (exact): d’ (W = –14, p = 0.3828), C (W = 4, p = 0.8438). (D) Cumulative distributions for hit and false alarm reaction times (RT) during mask and laser inhibition conditions. Curves pool trials from multiple sessions across 8 mice [Mask: n_hit_ = 2257, n_FA_ = 672; Laser: n_hit_ = 162, n_FA_ = 53]. Differences between distributions were tested with two-sample Kolmogorov-Smirnov tests (two-sided): Hits, D = 0.0427, p = 0.2955; False Alarms, D = 0.1676, p = 0.1921. (E) Hit and false alarm rates for mask control versus laser on post-sound trials (n = 8). Two-tailed Wilcoxon matched-pairs signed-rank test (exact): Hits (W = –28, p = 0.0547), FAs (W = 12, p = 0.4609). (F) Discrimination sensitivity and response bias values for mask control versus laser on post-sound trials (n = 8). Two-tailed Wilcoxon matched-pairs signed-rank test (exact): d’ (W = –28, p = 0.0547), C (W = 20, p = 0.1953). (G) Cumulative distributions for hit and false alarm reaction times (RT) during mask and laser inhibition conditions. Curves pool trials from multiple sessions across 8 mice [Mask: n_hit_ = 2356, n_FA_ = 88; Laser: n_hit_ = 767, n_FA_ = 30]. Differences between distributions were tested with two-sample Kolmogorov-Smirnov tests (two-sided): Hits, D = 0.0572, p = 0.0437; False Alarms, D = 0.1894, p = 0.3618. (H) Average reaction time by animal for hits (left) and false alarm (right) trials across mask and laser on during sound trials. One animal was excluded from the right panel because it had no false alarms in either condition. Two-tailed Wilcoxon matched-pairs signed-rank test (exact): Hits (n = 8, W = –22, p = 0.1484), FAs (n = 7, W = 16, p = 0.2188). (I) Average reaction time by animal for hits (left) and false alarm (right) trials across mask and laser on after sound trials. One animal was excluded from the right panel because it had no false alarms in the laser condition. Two-tailed Wilcoxon matched-pairs signed-rank test (exact): Hits (n = 8, W = –6, p = 0.7422), FAs (n = 7, W = –10, p = 0.4688).

**Figure S7.**
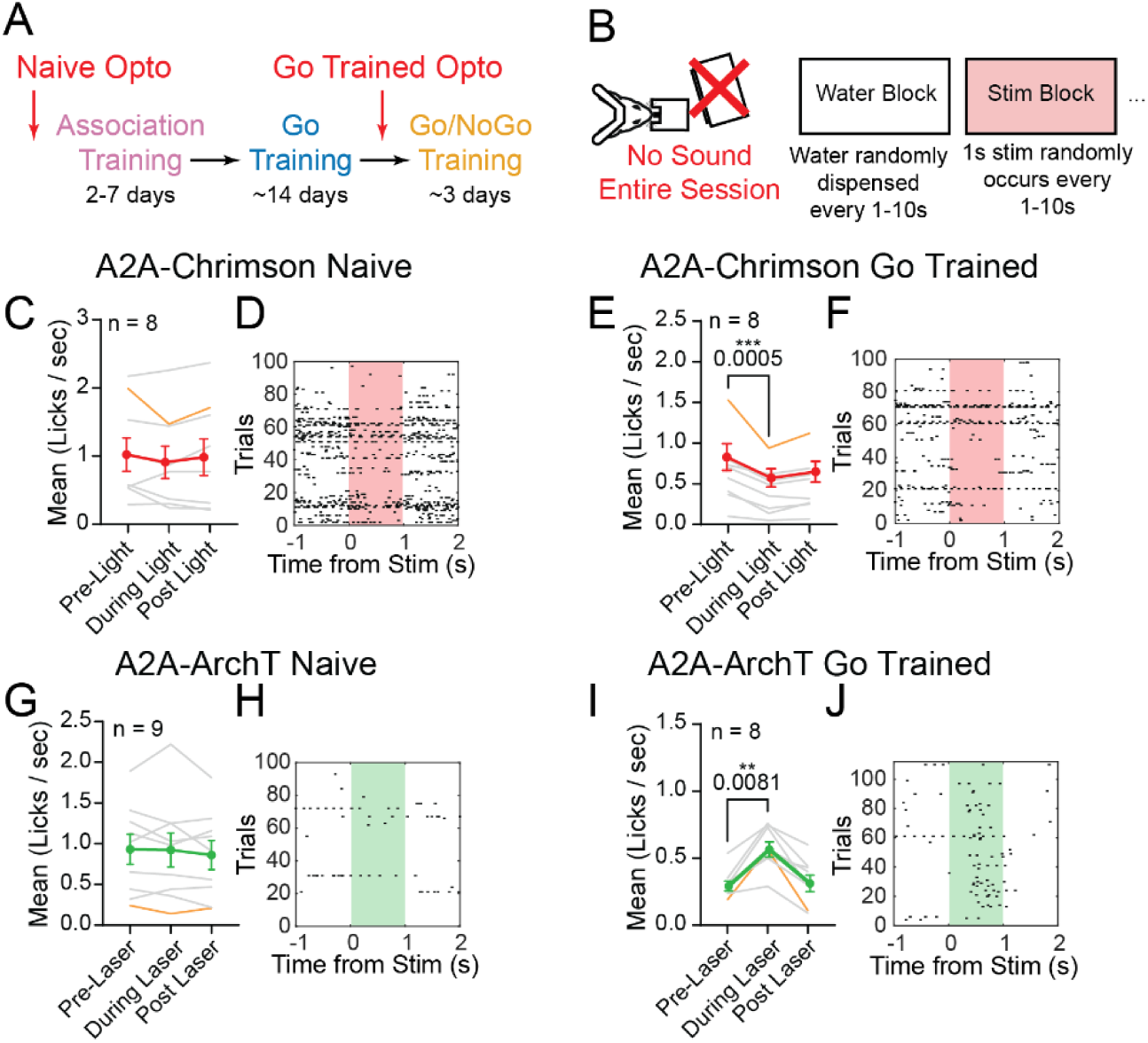
The indirect pathway can bias free licking probability only following training. (A) Training timeline with non-auditory opto session timepoints. (B) Schematic of the non-auditory optogenetic session. In these sessions, animals were encouraged to lick freely through random water deliveries presented in blocks alternating with stimulation periods. (C) Mean lick rates before, during and after optogenetic excitation of iSPNs in naïve mice. Gray lines signify individual animals with the red symbols representing mean ± SEM. The animal highlighted in orange displayed the greatest change in lick rate during go-trained stimulation (E) and was used to generate the data shown in D. Friedman RM ANOVA on ranks (3 conditions, n = 8): χ²(2) = 0.4516, p = 0.874). (D) Example lick raster for iSPN excitation in a naïve mouse before, during and after stimulation trials. The mouse selected displayed the greatest change in mean lick rate during go-trained stimulation. Individual licks denoted by black rasters with stimulation period represented by shaded region. (E) Mean lick rates before, during and after optogenetic excitation of iSPNs in go trained mice. Gray lines signify individual animals with the red symbols representing mean ± SEM. The animal highlighted in orange displayed the greatest change in lick rate during stimulation and was used to generate the data shown in F. Friedman RM ANOVA on ranks (3 conditions, n = 8): χ²(2) = 14.25, p < 0.0001****). Post hoc Dunn’s (adjusted p): Pre-Light vs During-Light p = 0.0005***, Pre-Light vs Post-Light p = 0.073, During-Light vs Post-Light p = 0.401. (F) Example lick raster for iSPN excitation in a go trained mouse before, during and after stimulation trials. The mouse selected displayed the greatest change in mean lick rate during go-trained stimulation. Individual licks denoted by black rasters with stimulation period represented by shaded region. (G) Mean lick rates before, during and after optogenetic inhibition of iSPNs in naïve mice. Gray lines signify individual animals with green symbols representing mean ± SEM. The animal highlighted in orange displayed the greatest change in lick rate during go-trained stimulation (I) and was used to generate the data shown in H. Friedman RM ANOVA on ranks (3 conditions, n = 9): χ²(2) = 2.667, p = 0.3285). (H) Example lick raster for iSPN inhibition in a naïve mouse before, during and after stimulation trials. The mouse selected displayed the greatest change in mean lick rate during go-trained stimulation. Individual licks denoted by black rasters with stimulation period represented by shaded region. (I) Mean lick rates before, during and after optogenetic inhibition of iSPNs in go trained mice. Gray lines signify individual animals with green symbols representing mean ± SEM. The animal highlighted in orange displayed the greatest change in lick rate during stimulation and was used to generate the data shown in J. Friedman RM ANOVA on ranks (3 conditions, n = 8): χ²(2) = 12, p = 0.0011**). Post hoc Dunn’s (adjusted p): Pre-Laser vs During-Laser p = 0.0081**, Pre-Laser vs Post-Laser p > 0.9999, During-Laser vs Post-Laser p = 0.0081**. (J) Example lick raster for iSPN inhibition in a go trained mouse before, during and after stimulation trials. The mouse selected displayed the greatest change in mean lick rate during go-trained stimulation. Individual licks denoted by black rasters with stimulation period represented by shaded region.

## Acknowledgements

This work was supported by the National Institute of Neurological Disorders and Stroke F31 fellowship (1F31NS130989-01) to S.F. and the National Institute of Mental Health R01 grant (R01MH136354) and Autism Spectrum Program of Excellence (ASPE) support to M.F. S.P. is supported as a Penn ASPE postdoctoral fellow.

We thank Felicia Davatolhagh and Edgar Diaz-Hernandez for their contributions to behavioral apparatus design, Nathan Volger for assistance with sound system calibration, and Allison Girasole for valuable discussions on animal training. We also thank Drs. Girasole and Long Ding for reading the manuscript.

